# DMSO derives Trophectoderm and Clonal Blastoid from Single Human Pluripotent Stem Cell

**DOI:** 10.1101/2023.12.31.573770

**Authors:** Samhan Alsolami, Arun Pandian Chandrasekaran, Yiqing Jin, Ismail M. Shakir, Yingzi Zhang, Gerardo Ramos-Mandujano, Baolei Yuan, Alfonso Saera-Vila, Juan Carlos Izpisua Belmonte, Mo Li

## Abstract

Human naïve pluripotent stem cells (nPSCs) can differentiate into extra-embryonic trophectoderm (TE), a critical step in the generation of the integrated embryo model termed blastoid. The current paradigm of blastoid generation necessitates the aggregation of dozens of nPSCs treated with multiple small molecule inhibitors, growth factors, or genetic modifications to initiate TE differentiation. The presence of complex crosstalk among pathways and cellular heterogeneity in these models complicates mechanistic study and genetic screens. Here, we show that a single small molecule, dimethyl sulfoxide (DMSO), potently induces TE differentiation in basal medium without pharmacological and genetic perturbations. DMSO enhances blastoid generation and, more importantly, is sufficient for blastoid generation by itself. DMSO blastoids resemble blastocysts in morphology and lineage composition. DMSO induces blastoid formation through PKC signaling and cell cycle regulation. Lastly, DMSO enables single nPSC-derived clonal blastoids, which could facilitate genetic screens for mechanistic understanding of human embryogenesis.

## INTRODUCTION

The trophectoderm (TE) plays a critical role in blastocyst development by forming a single-cell epithelial layer surrounding the inner cell mass (ICM) and regulating the movement of water to form a fluid-filled blastocoel. Comprehending the signaling requirements of TE is crucial for modeling the formation of human blastocysts, implantation, and placental development. In vitro derivation of TE cells is limited to early pluripotent stem cells (PSCs), such as naïve pluripotent stem cells (nPSCs)^1^. Recently, the ability to derive TE-like cells from nPSCs has fueled the generation of many blastocyst-like models termed blastoids ^2-8^. In vitro-derived TE cells can be produced by simultaneously inhibiting the ERK and TGFβ/Nodal pathways with PD0325901 (PD) and A83-01 (A83), respectively^1^. Brief induction of TE in so-called PXGL nPSCs using PD and A83 (PDA83) results in the formation of human blastoids, which consist of TE, primitive endoderm (PE), and epiblast (EPI)^4,6^. A multitude of other inhibitors and growth factors, including VPA (HDAC inhibitor)^3,5,7,8^, CHIR99021 (GSK3 inhibitor), BMP4^2,3,8,9^, hEGF^5,7^, IM-12 (GSK3b inhibitor)^5,7^, and WH- 4-023 (LCK/SRC inhibitor)^5,7^ are purported to facilitate TE differentiation and/or blastoid formation. Although these external interventions are successful in inducing TE from nPSCs to different extent, they by no means represent the endogenous mechanisms underlying the first lineage segregation event in humans where the zygote can self-organize the lineage differentiation program. These factors target multiple pathways that have complex crosstalk among them, which makes it challenging to understand the logic of the TE program. Complex culture conditions involving many inhibitors and growth factors often require labor-intensive optimization to accommodate cell lines of different genetic background and the use of supra-physiological levels of signaling manipulation has been associated with limited developmental potential^6,10^. Additionally, existing protocols have relied on aggregates of human PSCs for blastoid generation. The variability among cellular aggregates hampers culture reproducibility and forward genetic screens to understand the gene regulatory networks in human embryogenesis. In this study, we demonstrate that a single molecule, DMSO, without previously used inhibitors, growth factors, or genetic manipulations^3^, can induce TE from human nPSCs and promote the generation of human blastoids from single nPSCs.

## RESULTS

### DMSO promotes the exit of pluripotency and trophectoderm differentiation of human naive PSCs

In vivo, the mechanisms underlying the first lineage segregation event, where the totipotent blastomeres differentiate into an outer TE layer and an ICM shortly after the morula stage, remain largely unknown. Hippo pathway effectors^11-13^, GATA transcription factors (TFs)^1^, and hydraulic fracturing of cellular contacts^14^ have been proposed to underlie TE lineage specification in the blastocyst. Cell surface fluctuations have been shown to regulate early embryonic lineage sorting^15^. DMSO is known to downregulate pluripotency genes, which promotes differentiation in human primed embryonic stem cells (ESCs)^16,17,18^. Its effects on plasma membrane fluidity and permeability have been well documented and are suggested to induce cellular differentiation and fusion^19-23^. We hypothesized that DMSO could bias nPSCs toward the TE branch of the first cell fate bifurcation.

To test this hypothesis, we subjected nPSCs to DMSO treatment in N2B27 basal medium in two-dimensional (2D) culture (Fig. 1a). Remarkably, in 3-4 days, DMSO-treated nPSCs differentiated into numerous cystic structures, which was reminiscent to the TE induction by PDA83 reported previously^1^. DMSO also further enhanced the effect of PDA83 on epithelial cyst formation, resulting in a confluent layer of fluid-filled cysts (Fig. 1a). The highly reproducible (>15 trials) formation of cystic structures suggested that DMSO promoted differentiation of functional nPSC-derived TE-like cells. Consistently, DMSO treatment downregulated the pluripotency marker SOX2 and activated key TE TF GATA3 in the majority of cells (Fig. 1b, Supplementary Fig. S1a, b). SOX2 was expressed in 98.3±1.1% of cells cultured in N2B27, whereas it was present in only about 3.7±1.3% of DMSO treated cells and about 39.5±11.1% in PDA83 treated cells (Fig. 1c). Conversely, GATA3 was absent in N2B27 cultured cells but expressed in 69.5±11.1% of DMSO treated cells and 53.8±6.4% of PDA83 treated cells (Fig. 1c). Three-dimensional (3D) imaging reconstruction showed that the lining of the cysts consisted of a single layer of cells with strong nuclear GATA3 staining and prominent cortical F-actin at the boundaries of cells, consistent with a polarized TE-like epithelium (Fig. 1d, Supplementary Fig. S1c). Flow cytometry analysis unveiled that DMSO upregulated the TE marker TROP2 (encoded by the *TACSTD2* gene) more potently than PDA83 and worked additively with PDA83 to promote the TE fate in most cells (Fig. 1e, Supplementary Fig. S1d,e). Gene expression analysis by quantitative reverse transcription PCR (qRT-PCR) further corroborated the sharp downregulation of naïve pluripotency genes (*NANOG* and *KLF4*) and upregulation of TE marker genes (*GATA3*, *ADAP2*, and *TACSTD2*/*TROP2*) in DMSO treated cells, which also prevented the upregulation of primed pluripotency markers (*ZIC2* and THY1/CD90 by flow cytometry, Fig. 1f, Supplementary Fig. S1e,f).

**Fig. 1.**
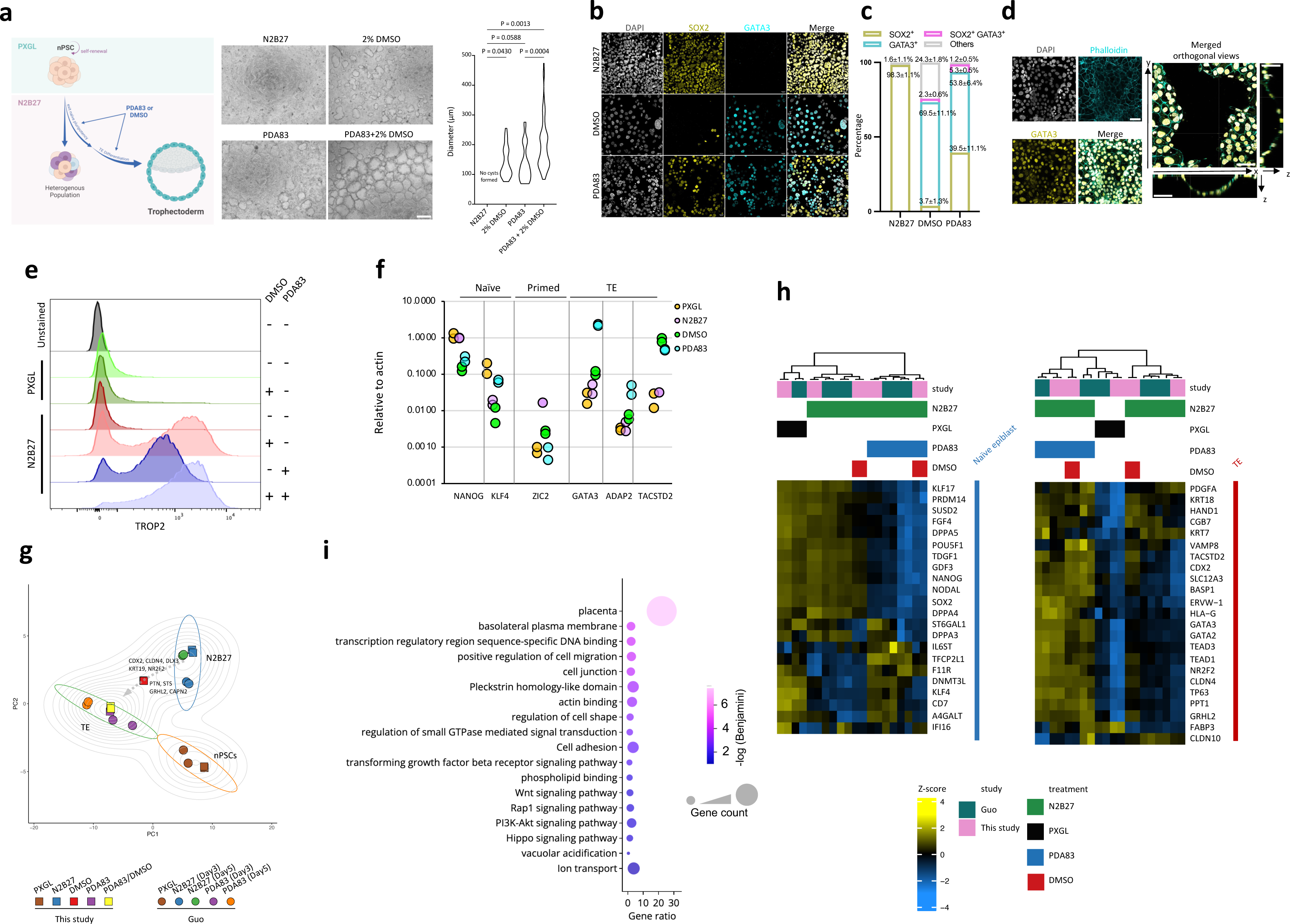
DMSO drives trophectoderm differentiation. **a**, Schematic depicting the DMSO function in promoting TE fate and in inhibiting primed PSC formation (**left**). Representative brightfield images of the day 4 structures show the effect of cyst formation in different conditions (**middle**). Scale bar, 400 μm. The violin plot shows the diameter of the cysts formed (**right**). Data are presented as the mean and standard deviation. N2B27: no cysts; 2% DMSO: n = 26 cysts; PDA83: n = 31 cysts; and PDA83+2% DMSO: n = 84 cysts. One-way ANOVA followed by the Tukey post hoc test was used, and the P values are as indicated. **b**, Immunofluorescence analysis shows the expression of SOX2 (yellow) and GATA3 (cyan) in N2B27, 2% DMSO, and PDA83 conditions. Scale bar, 20 μm. **c**, Quantification of immunofluorescence data from (**b**) identifies four distinct cell types: SOX2-positive cells, GATA3-positive cells, cells co-expressing SOX2 and GATA3, and other cells (n = 2). **d**, Immunofluorescence analysis shows the expression of Phalloidin (cyan) and GATA3 (yellow) in 2% DMSO condition (**left**). The orthogonal view demonstrates 2D trophectoderm sphere formation (**right**). Scale bar, 50 μm. **e**, FACS histogram shows the TROP2 expression in various conditions (n = 3). **f**, qRT-PCR shows the expression pattern of various genes in naive, primed, and TE states under different culture conditions (n = 2). **g-i**, PCA plot of integrated bulk RNAseq analysis of nPSC, N2B27, and TE differentiation from this study and previously published data. The dotted arrow indicates the top PCA loading genes (*CDX2*, *CLDN4*, *DLX3*, *KRT19*, *NR2F2*, *PTN*, *STS*, *GRHL2*, and *CAPN2*) between the N2B27 and DMSO samples. (**g**), Heatmap shows the differential expression pattern of genes in different treatments (**h**). GO analysis of differentially expressed genes between DMSO and N2B27 conditions (**i**).

To gain a comprehensive view of the DMSO gene transcription program, we performed RNA sequencing (RNA-seq) of nPSCs in PXGL and various differentiation conditions. Principal component analysis (PCA) showed excellent reproducibility of the experiments and agreement between our data and previously published datasets^1^ of nPSC cultured in PXGL medium, N2B27 basal medium, and N2B27 with PDA83 (Fig. 1g). The DMSO plus PDA83 samples closely clustered with TE differentiated samples (PDA83) and away from the undifferentiated (nPSC/PXGL) and heterogeneously differentiated samples (N2B27) in the PCA plot (Fig. 1g). Interestingly, DMSO treatment alone induced a differentiation trajectory in nPSCs closely aligning with the TE trajectory in both PC1 and PC2, as shown in (Fig. 1g); PCA gene loading analysis further reveals that DMSO PCA localization is predominantly influenced by TE related genes such as *CDX2*, *CLDN4*, *DLX3*, *KRT19*, and *NR2F2* (Fig. 1g). DMSO with or without PDA83 coordinately induced TE transcripts (e.g., *TACSTD2*, *GRHL2*, *TEAD1*, *CDX2*, *KRT18*, and *CGB7*) and downregulated naïve transcripts (e.g., *NANOG*, *KLF17*, *KLF4*, *DNMT3L*, and *SUSD2*) (Fig. 1h, Supplementary Fig. S1e). Differentially expressed genes between DMSO-directed differentiation and random differentiation in N2B27 basal medium were significantly enriched in gene ontology (GO) terms consistent with TE cell fate, including the tissue term placenta, cellular component terms basolateral plasma membrane and cell junction, and biological process germs cell adhesion, ion transport, and Hippo signaling (Fig. 1i). Together these data suggested that a single small molecule DMSO could induce differentiation of TE-like cells from nPSCs without the need for inhibitors of signaling pathways or genetic manipulation.

### DMSO directs chromatin accessibility changes that promote TE differentiation

To further understand the cis-regulatory elements underlying the transcriptomic changes induced by DMSO, we performed an assay for transposase-accessible chromatin with sequencing (ATAC-Seq) in PXGL, N2B27, PDA83, PDA83+DMSO, and DMSO samples. The chromatin accessibility landscape of DMSO treated samples (DMSO alone and PDA83+DMSO) clustered close to that of PDA83 differentiated TE cells (Fig. 2a). DMSO increased chromatin accessibility in the neighborhood of TE genes (e.g., *GATA3*, *NR2F2*, *TEAD1*, *TACSTD2*, and *TP63*), a chromatin dynamics profile shared with PDA83 (Fig. 2b,c). Notably, DMSO led to diminished ATAC-seq peaks in many naive (e.g., *KLF17*, *SUSD2*, and *DNMT3L*) and primed (e.g., *ZIC2*, *OTX2*, and *THY1*) pluripotency markers, which were reminiscent of ATAC-seq peaks of PDA83 and corroborated with gene expression changes induced by DMSO (Fig. 2c, Fig. 1f,h, and Supplementary Fig. S1b).

**Fig. 2.**
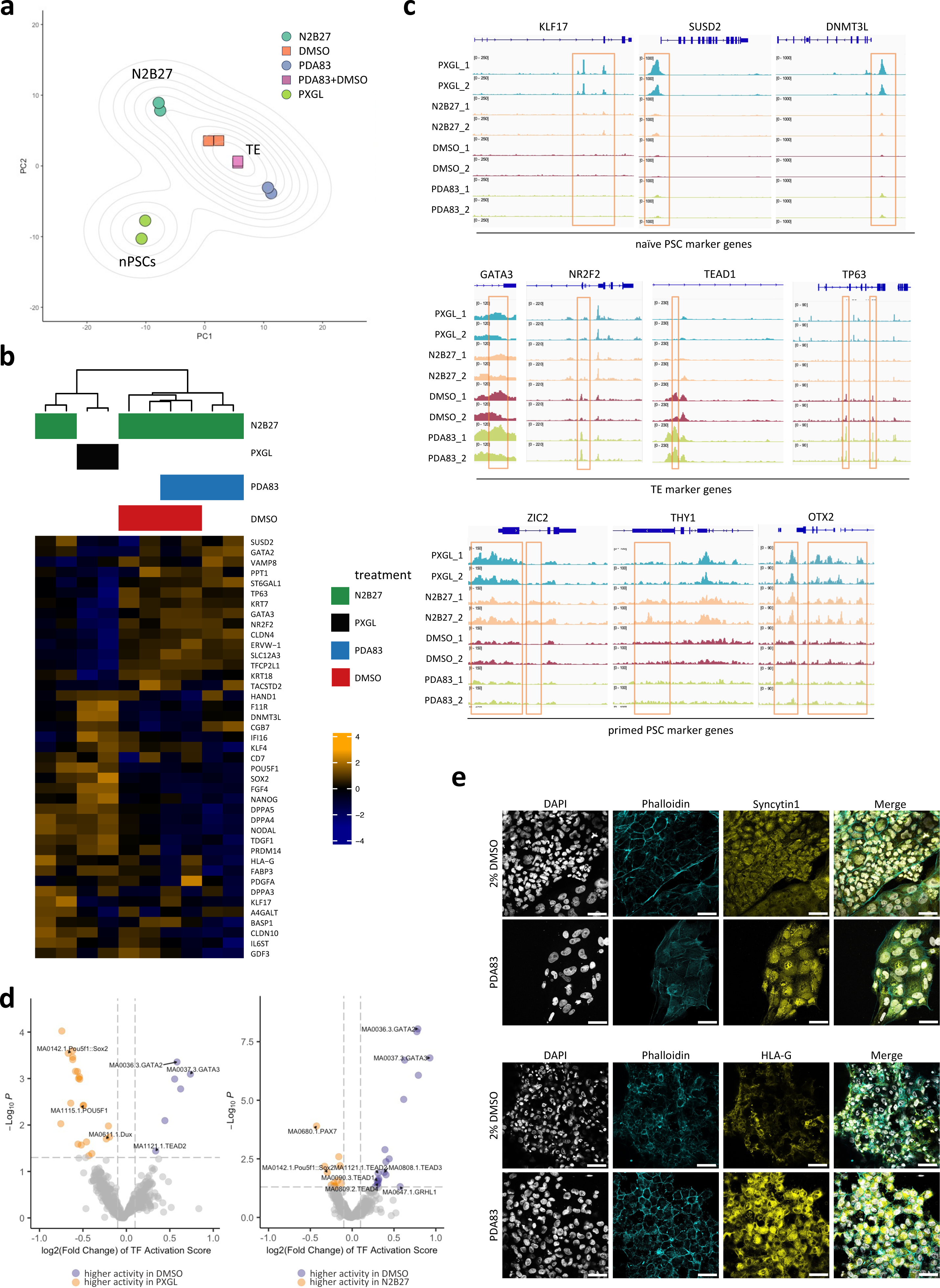
ATAC-seq analysis reveals the role of DMSO in TE fate. **a-b**, PCA (**a**), and heatmap (**b**) depict open chromatin variations in different treatment conditions. **c**, IGV plots of markers for naïve PSCs (**top**), trophectoderm (**middle**), and primed PSCs (**bottom**). **d**, The volcano plot shows the comparisons of transcription factor activity across the epigenome of DMSO vs. PXGL (**left**) and DMSO vs. N2B27 (**right**) conditions. **e**, Immunofluorescence analysis shows the expression pattern of Phalloidin (cyan) vs. Syncytin1 (yellow) (**top**) and Phalloidin (cyan) vs. HLA-G (yellow) (**bottom**) under DMSO and PDA83 conditions (n = 3). Scale bar, 50 μm.

Moreover, motif enrichment analysis showed that DMSO had significantly higher activation scores^24^ of TE TFs (e.g., *GATA2*, *GATA3*, *TEAD2*, and *TEAD3*) and lower activation scores of naïve TFs (e.g., *POU5F1*, *SOX2*, and *Dux*) (Fig. 2d, Supplementary Fig. S2a-c). Finally, DMSO-induced TE-like cells could be further differentiated into syncytin-1/ERVW-1-positive syncytiotrophoblasts (STB) and HLA-G-positive extravillous trophoblasts (EVT) using established protocols^2,25^ (Fig. 2e). Together, these data show that DMSO alone can induce a TE-like gene expression and cis-regulatory programs consistent with the differentiation of trophectoderm from a naive pluripotency state.

### DMSO enhances pre- and peri-implantation blastoid formation

Since DMSO dramatically enhanced TE differentiation efficiency (Fig. 1a,e, Supplementary Fig. S1e) of nPSCs induced by PDA83, we hypothesized that it could facilitate human blastoid formation using a method^6^ that relies on PDA83, the Hippo pathway inhibitor lysophosphatidic acid (LPA), leukemia inhibitory factor (LIF), and Y-27632 (a ROCK inhibitor) (hereafter referred to as PALLY). We first compared the timing and dosage of DMSO treatment in the PALLY protocol. Interestingly, DMSO significantly promoted the cavitation process of the PALLY condition, as evidenced by the larger diameter of the cavitated structures (Supplementary Fig. S3a). The average diameter of cavitated structures correlated positively with DMSO dosage from 0.5–2% (Supplementary Fig. S3a, right panel), while 0.1% DMSO had little effect (data not shown). We, therefore, decided to use 0.5% DMSO (Fig. 3a), which produced diameters consistent with human blastocysts 5-6 days after fertilization^26,27^, while maintaining high quality cavitated structures consistently (Supplementary Fig. S3b,c). The cavitated structures in PALLY+DMSO were morphologically indistinguishable from PALLY blastoids and underwent the same morphogenic process of compaction, cavitation, and expansion (Supplementary Movies 1 and 2 and Supplementary Fig. S3c). Immunofluorescence showed that, like PALLY blastoids, the cavitated structures in PALLY+DMSO contained a single outer epithelial layer of TE-like cells (positive for GATA3 and CK18), a cluster of EPI-like cells (positive for SOX2 and OCT4), and an arrangement of PE-like cells on the cavity-facing side of the inner cell cluster (positive for GATA4) (Supplementary Fig. S3d), therefore proving their blastoid identity.

**Fig. 3.**
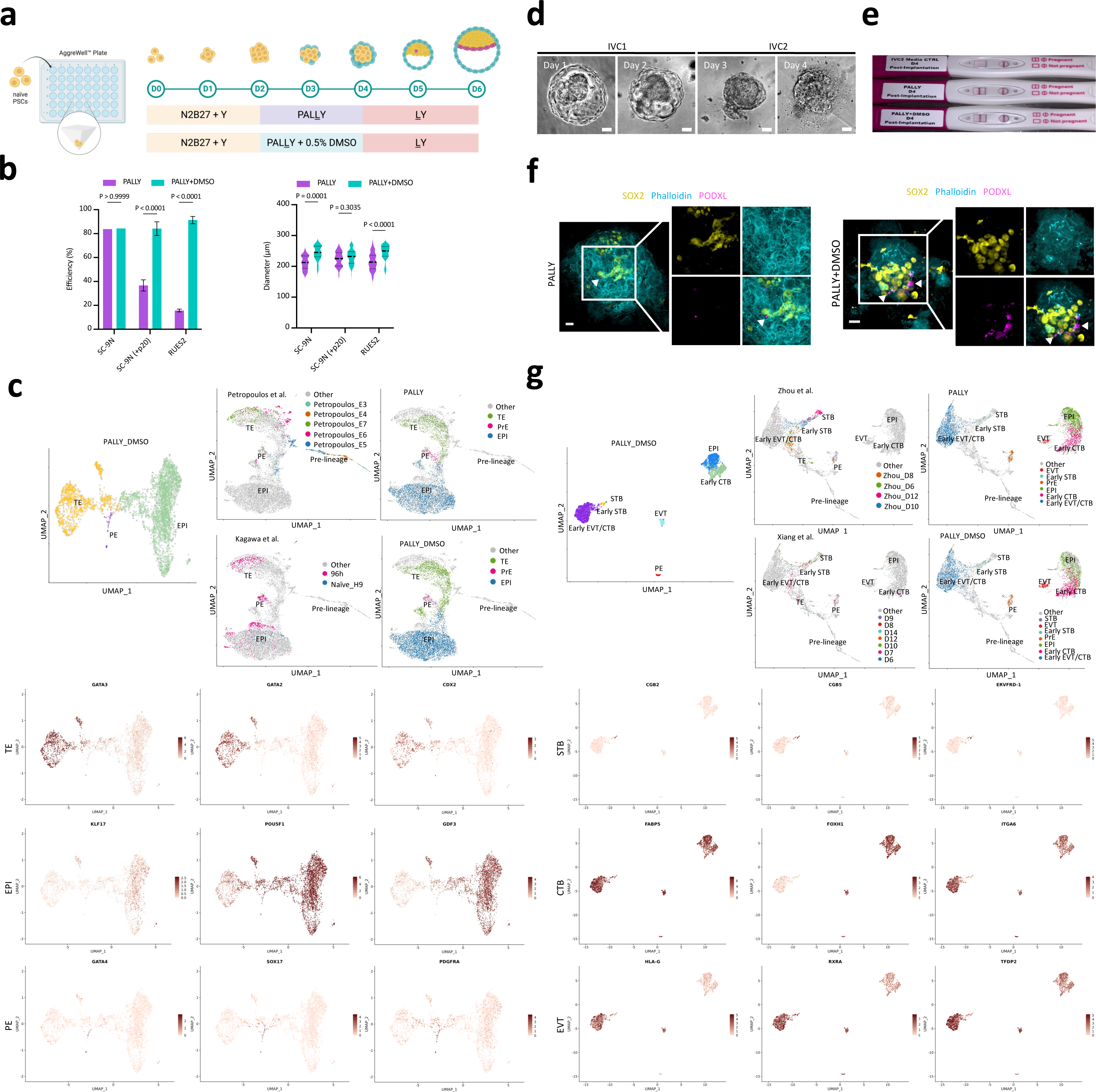
Synergistic effect of DMSO in PALLY conditions improves blastoid generation. **a**, Schematic depicts the DMSO dosage in PALLY condition. **b**, Comparative analyses of efficiency (**left**), and size (**right**) of the blastoids in different cell lines. SC-9N (+p20) represents 20 passages after chemically resetting the SC-9N line. Data are presented as the mean and standard deviation. Two-way ANOVA followed by Bonferroni’s post hoc test was used, and the P values are as indicated. **c**, UMAP of the transcriptome of 3244 single cells of the pre-implantation PALLY+DMSO blastoids with cell type annotation (**top left panel**). UMAP projection of integrated datasets showing cells from this study and published studies by Kagawa et al. and Petropoulos et al. (**top right panels**). Feature plots of markers of each blastocyst lineage (TE, EPI, and PE; **bottom panels**). **d**, Brightfield images show the successful implantation of blastoid in PALLY+0.5% DMSO condition. Scale bar, 50 μm. **e**, Commercial pregnancy test kit detects stimulated CGý under PALLY+0.5% DMSO condition. **f**, Immunofluorescence analysis shows the expression of SOX2 (yellow), Phalloidin (cyan), and PODXL (magenta) in PALLY+0.5% DMSO condition. Scale bar, 60 μm. **g**, UMAP of the transcriptome of 3875 single cells of the attached PALLY+DMSO blastoids with cell type annotation (**top left panel**) and integrated datasets showing cells from this study and previously reported conditions (**top right panels**). Feature plots of markers of each blastocyst lineage (STB, CTB, and EVT; **bottom panels**).

We next examined the effect of PALLY and PALLY+DMSO on different nPSC lines and nPSCs of different passages. We found significant variabilities in the efficiency of blastoid formation under the PALLY condition between different nPSC lines and the same nPSC line of different passages. Importantly, adding DMSO to PALLY successfully mitigated the inconsistent cavity formation observed across the samples (Fig. 3b). Thus, the above findings unambiguously demonstrate the utility of DMSO in achieving consistent and reliable blastoid formation under the PALLY condition.

Single-cell RNA-seq analysis revealed that blastoids from both PALLY+DMSO and PALLY conditions possessed well-delineated cellular populations expressing marker genes^1,6,28^ specific to the three lineages of the blastocyst–TE (e.g., *GATA3*, *GATA2*, and *CDX2*), EPI (e.g., *POU5F1*, *KLF17*, and *GDF3*), and PE (e.g., *GATA4*, *SOX17*, and *PDGFRA*) (Fig. 3c, Supplementary Fig. S3e, and Supplementary table 2). The transcriptomic profiles of cells in PALLY+DMSO blastoids showed good overlap with those in human blastocysts^28^ and PALLY blastoids (both this study and a published study^6^) but not with human embryos prior to lineage segregation, which is consistent with the expected developmental stage of the blastoids (Fig. 3c, Supplementary Fig. S3e). To test the specification of cell fates in blastoids functionally, we re-derived nPSCs (OCT4^+^NANOG^+^) and trophoblast stem cells (GATA3^+^CK7^+^) from PALLY+DMSO and PALLY blastoids as previously described^6^ (Supplementary Fig. S3f).

After expansion and hatching, the blastocyst implants into the endometrium through TE, subsequently giving rise to cytotrophoblasts (CTBs), STBs, and EVTs to support embryo development. To further study if blastoids could model the peri-implantation process, they were placed on Matrigel pre-coated dishes and cultured for four days (see *Methods*). PALLY-DMSO blastoids collapsed, and the outer TE invaded the matrix, while the inner cells differentiated into an epithelial-like disc (Fig. 3d). The TE derivatives (CK18^+^GATA3^+^phalloidin^+^) spread outward from attached blastoids and secreted human chorionic gonadotropin hormone (hCG) into the medium above the positive detection threshold of commercial pregnancy tests (Fig. 3e, Supplementary Fig. S3g). Post-implantation EPI goes through epithelialization and another cavitation event to form the amnion and the amniotic cavity. Interestingly, the disc-like structure contained well-packed SOX2^+^ EPI analogs with strong phalloidin and PODXL staining in between cells on the periphery of the colony, suggesting the formation of an early amniotic cavity-like structure. In contrast, PODXL^+^ signals were only sparsely found in attached PALLY blastoids at day 4 of attachment (Fig. 3f). GATA4^+^ PE derivatives were also present near the periphery of the SOX2^+^ colony (Supplementary Fig. S3g).

Furthermore, scRNAseq showed that attached PALLY+DMSO blastoids (PALLY blastoids as well) contained well-delineated cellular populations expressing marker genes^1,6,28^ specific to STB (e.g., *CGB2*, *CGB5*, and *ERVFRD-1*), CTB (e.g., *FABP5*, *ITGA6*, and *FOXH1*), and EVT (e.g., *HLA-G*, *RXRA*, and *TFDP2*) (Fig. 3g, Supplementary Fig. S3h). Cells in attached blastoids aligned well transcriptionally with cells in human post-implantation embryos^27,29^ but not pre-implantation blastocysts^28^, which is consistent with their expected developmental stage (Fig. 3g, Supplementary Fig. S3h, and Supplementary table 2). Together, these observations demonstrate that by promoting TE differentiation, DMSO facilitates the efficient formation of blastoids that can differentiate into lineages analogous to those found in post-implantation embryos.

### DMSO alone induces blastoids without lineage instructive cues

Given the fact that DMSO induces the TE lineage (Fig. 1) and improves PALLY blastoid formation (Fig. 3), we sought to investigate whether DMSO alone could induce blastoid formation in nPSCs. Since higher concentrations of DMSO at the beginning of the PALLY treatment dramatically promoted cavitation (Supplementary Fig. S3a), we treated human nPSCs with 2% DMSO to evaluate blastoid formation (Fig. 4a). Upon exposure to 2% DMSO in N2B27 medium, blastocyst-like cavitated structures formed on day 6- 7 (D6-7, Fig. 4b), while the efficiency of cavitation was relatively low (Supplementary Fig. S4a). DMSO is known to slow down cell cycle progression^30,31^(also see discussion in the next section). Consistently, we noted that D7 structures were of better quality and consistency (Supplementary Fig. S4b-e). Immunofluorescence analysis of D7 blastoids revealed the presence of three founding lineages (GATA3^+^, SOX2^+^, and GATA4^+^ cells, Fig. 4c). Recent studies have shown that LIF and LPA can improve human blastoid development^6,32^. We then tested whether LIF and LPA could improve the efficiency of DMSO-derived blastoid formation. The addition of LIF and LPA, either separately or in combination, to the DMSO protocol resulted in the formation of cavitated structures (Supplementary Fig. S4f,g). Interestingly, adding LPA at the end of the differentiation window (conditions #1, #5, and #7) significantly increased blastoid cavitation efficiency (Fig. 4d and Supplementary Fig. S4h) without affecting blastoid size (Supplementary Fig. S4i). Adding LIF alone with DMSO did not increase the number of cavitated structures (condition #2). Immunofluorescence analysis showed that the combination of LPA and DMSO resulted in the differentiation of all three lineages, while LIF was dispensable (Supplementary Fig. S4j,k). These results suggest that Hippo inhibition facilitates the second lineage segregation in DMSO-induced blastoid generation.

**Fig. 4.**
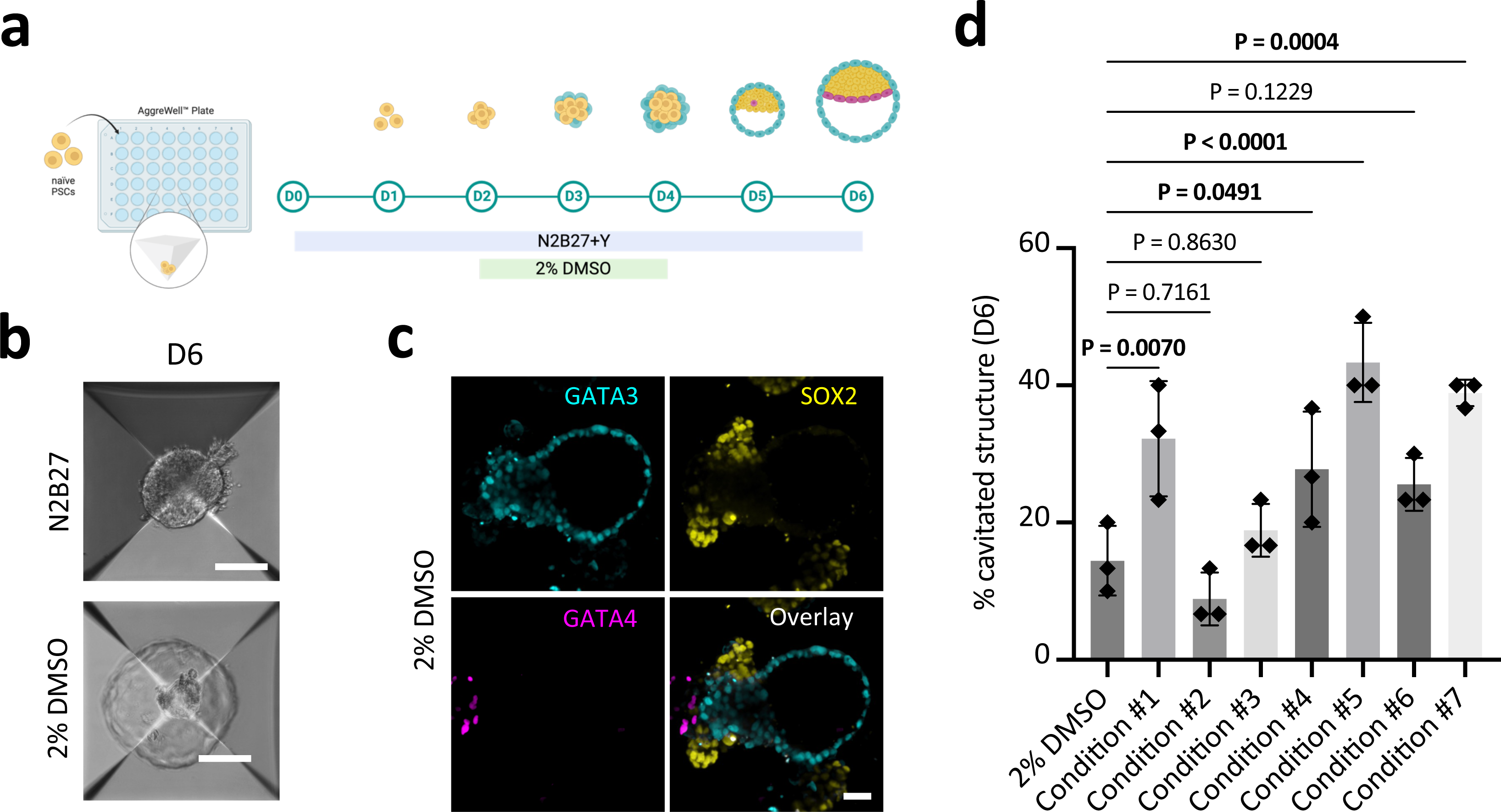
Derivation of human blastocyst-like structures from DMSO-treated nPSCs. **a**, Schematic representation of the generation of blastocyst-like structures using only DMSO. **b**, Representative phase-contrast images of DMSO-derived cavitated structures harvested from the microwells on D6. Scale bar, 100 μm. **c**, Immunofluorescence analysis of the TE marker (GATA3; cyan), EPI marker (SOX2; yellow), and PE marker (GATA4; magenta) of 2% DMSO-derived structures. Scale bar, 50 μm. **d**, Graph shows the percentage cavitation in different treatment conditions in D6 structures. Data are presented as the mean and standard deviation of three independent experiments. One-way ANOVA followed by the Dunnett post hoc test was used, and the P values are as indicated.

### Lineage specifying roles of DMSO require aPKC

We next sought to elucidate how DMSO promotes TE and blastoid formation. DMSO has a wide range of bioactive properties, including acting as cryoprotectant^33^, a solvent and penetration enhancer^34^, and a potent inducer of cellular differentiation ^19-23^. However, its mode of action at the cellular level remains unclear. We demonstrated that DMSO significantly accelerates cavitation and subsequent blastoid development of nPSCs. Cavitation, a critical event in embryogenesis, requires the differentiation of a polarized fluid-tight TE epithelia, which is actuated by Hippo signaling pathway effectors (e.g., nuclear *YAP1*) and TE-specific TFs (e.g., *GATA3*) in humans^1,11,13^. Atypical protein kinase C (aPKC) has been suggested to be the upstream regulator that initiates the TE program by setting up apical-basal polarity in the outer cells of the morula^12^. We, therefore, hypothesized that aPKC activity is required in DMSO-induced cavitation. Immunofluorescence revealed that aPKC was initially expressed in all cells of day 3 aggregates of nPSCs cultured in N2B27 basal media containing DMSO. At day 6, aPKC became highly restricted to TE cells (GATA3^+^) in both DMSO and PDA83 conditions (Fig. 5a, Supplementary Fig. S5a-d). Consistently, several aPKC-regulated cell polarity genes (e.g., *AMOT*, *GRHL2*, *PARD3B*, *PARD6A/B*, *RAB25*, *YAP1*, etc.) were induced by DMSO treatment (Supplementary Fig. S5e). Notably, aPKC colocalizes with TE but not ICM (OCT4^+^) cells (Fig. 5b).

**Fig. 5.**
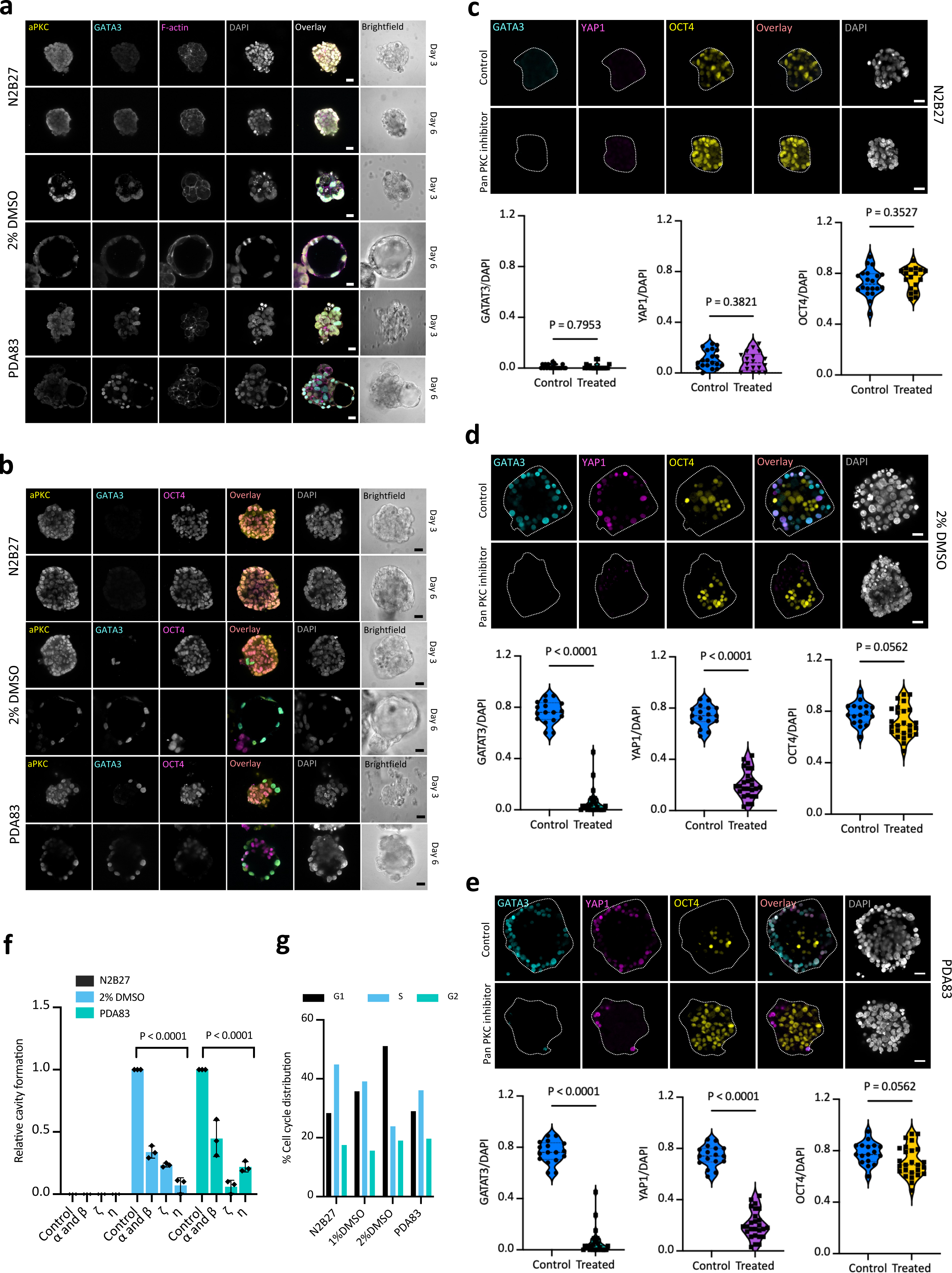
DMSO is a downstream regulator of the aPKC pathway. **a**, Representative immunofluorescence images showing the expression of aPKC (yellow), GATA3 (cyan), and F-actin (magenta) of D3 and D6 structures in N2B27, 2% DMSO, and PDA83 conditions. Scale bar, 25 μm. **b**, Representative immunofluorescence images showing the expression of aPKC (yellow), GATA3 (cyan), and OCT4 (magenta) of D3 and D6 structures in N2B27, 2% DMSO, and PDA83 conditions. Scale bar, 25 μm. **c-e**, Representative immunofluorescence images of D6 structures treated with pan PKC inhibitor showing the expression of GATA3 (cyan), YAP1 (magenta), and OCT4 (yellow) in N2B27 (**c**), 2% DMSO (**d**), and PDA83 (**e**) conditions. Scale bar, 25 μm. Quantification of GATA3, YAP1, and OCT4 normalized fluorescence intensity of the structures in the above conditions are provided in the respective panel. A two-tailed t-test was used, and the P values are as indicated. **f**, Relative blastoid generation efficiency after treatments with different inhibitors of PKC isoforms in N2B27, 2% DMSO, and PDA83 conditions. Data are presented as the mean and standard deviation of three independent experiments. Two-way ANOVA followed by Tukey’s post hoc test was used, and the P values are as indicated. **g**, Bar graph shows the percentage distribution of the cell cycle in N2B27, 1% DMSO, 2% DMSO, and PDA83 conditions.

Next, to test the requirement of PKC activity in DMSO-induced cavitation, we used a pan-PKC inhibitor Gö6983 that has been shown to block cavitation in human blastoids^5^ in a 2D TE differentiation assay. Epithelial cyst formation in either DMSO or PDA83 condition was severely abolished by Gö6983 (Supplementary Fig. S5f). Furthermore, Gö6983 inhibited the expression of GATA3 and YAP1 but not that of OCT4 (a marker for ICM) in 3D culture (Fig. 5c-e).

To study the role of specific PKC isoforms in cavitation, we used isozyme peptide inhibitors specific to PKCα/β, PKCζ, and PKCη. PKCη and PKCζ inhibitors significantly blocked cavitation and blastoid formation in DMSO and PDA83 treatment conditions, compared to the N2B27 control (Fig. 5f, Supplementary Fig. S5g). Our findings collectively demonstrate that PKC signaling is essential for DMSO-induced blastoid formation. These results provide new insights into the mechanisms by which DMSO stimulates TE differentiation and cavitation.

Retinoic acid (RA), like DMSO, is known as a differentiation inducer in leukemia cells^35^. RA signaling is essential for the totipotency program in mouse ESCs and zygotic genome activation in mammalian development^36^, and it is also required for trophoblast formation and ICM differentiation in porcine blastocysts^37^. Interestingly, our ATAC-seq analysis showed that the motifs of the nuclear receptors RARA/B/G (retinoic acid receptors alpha, beta, and gamma) and RXRA/B/G (retinoid X receptors alpha, beta, and gamma) were significantly enriched in the DMSO samples (Supplementary Fig. S2b,c). Thus, we investigate whether the effects of DMSO are mediated through RA signaling by treated nPSCs with RA in 2D culture for four days. In contrast to DMSO, RA did not induce cyst formation (Supplementary Fig. S5h). Consistently, FACS analysis revealed that RA was far less potent in the differentiation of TROP2^+^ TE-like cells (Supplementary Fig. S5i). These results suggest that RA signaling is unlikely to underline DMSO- induced blastoid formation.

We noticed that many cell cycle-related genes were coordinately downregulated in the DMSO condition (Supplementary Fig. S5j), which is consistent with published studies^30,31^. Cell cycle analysis by FACS showed that DMSO caused an accumulation of cells in the G1 phase, suggesting DMSO prolonged G1 (Fig. 5g). Phosphorylation of the retinoblastoma (Rb) protein is required for G1/S transition^38^. Western blot analysis revealed that DMSO treatment significantly decreased the fraction of phosphorylated Rb (pRb-Ser480) compared to the N2B27 control (Supplementary Fig. S5m), which coincided with epithelial cyst formation and TE marker TROP2 induction (Supplementary Fig. S5k and S5l). These results suggest that the mechanism underlying DMSO-induced TE differentiation is likely in part mediated by Rb and cell cycle regulation, which is reminiscent of the differentiation-promoting effect of DMSO in primed PSCs^30^.

### DMSO promotes clonal blastoid formation from a single nPSC

Current blastoid models recapitulate the sequence and timing of blastocyst development^2-6,8,39^. However, none of them have been shown to derive blastoids from a single cell, even though human blastocysts develop from a single cell–the zygote. To mimic the human blastocyst development, we attempted to generate clonal blastoids from single nPSCs. To achieve this, we seeded single human nPSCs at an appropriate density. A detailed protocol is described in *Methods*, and a schematic is depicted in Supplementary Fig. S6. A single nPSC was allowed to grow for 6-8 days, followed by PALLY and LY treatments for 48 hours each (Fig. 6a). Upon exposure to PALLY, cavitated structures were formed (Fig. 6b). Immunofluorescence analysis confirmed the presence of trilineage structures (GATA3^+^ for TE, SOX2^+^ for EPI, and GATA4^+^ for PE) (Fig. 6c), which is comparable to blastoids from a heterogeneous aggregate of 50-100 cells. Furthermore, supplementing DMSO to PALLY treatment resulted in cavitated structures (Fig. 6b) and trilineage populations (Fig. 6c) with a significantly increased efficiency (Fig. 6d), further demonstrating the indispensable role of DMSO for efficient blastoid formation.

**Fig. 6.**
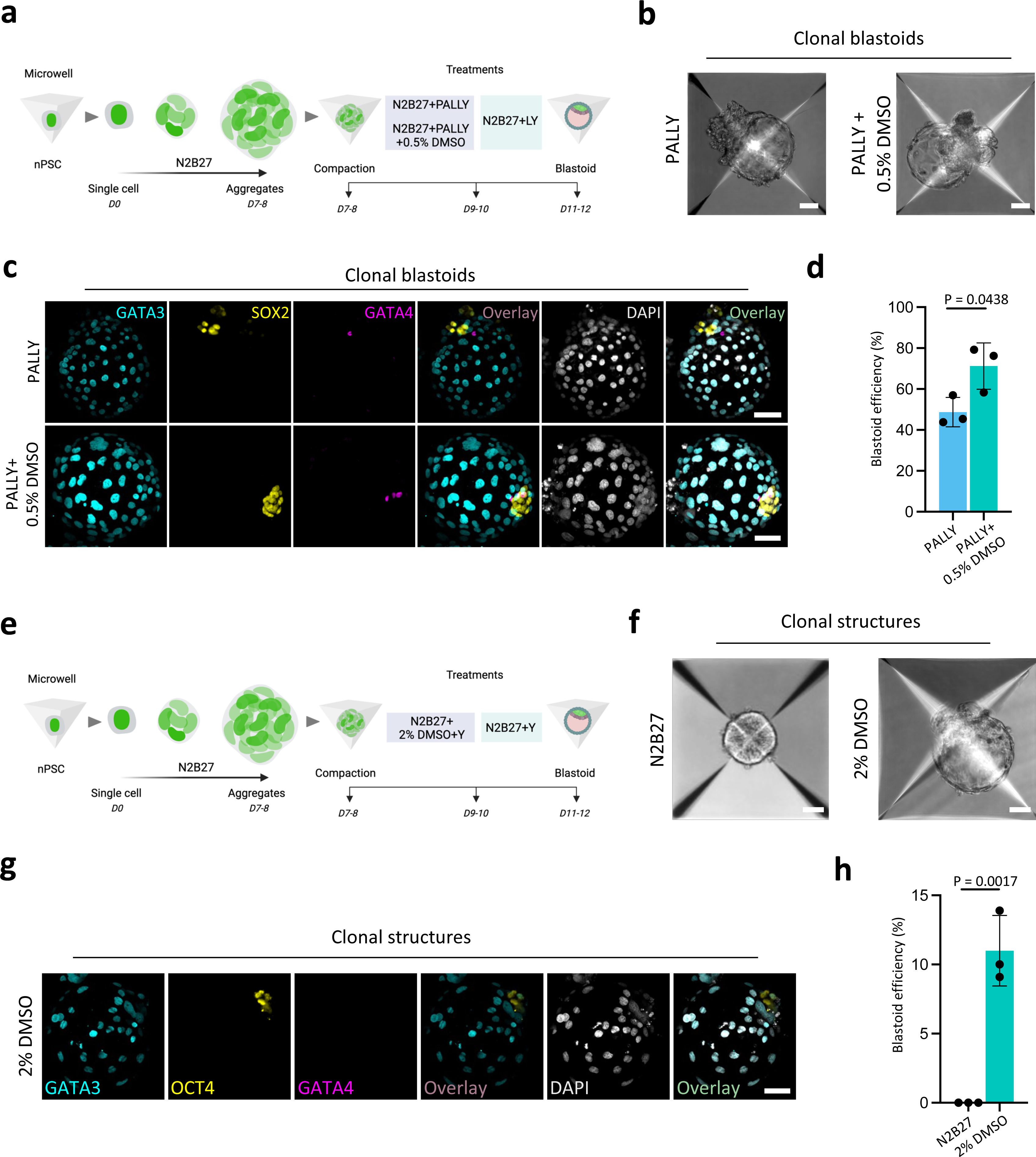
Clonal blastoid derivation from a single nPSC. **a**, Schematic depicting the generation of clonal blastoids in PALLY and PALLY+0.5% DMSO conditions. **b**, Representative phase-contrast images showing the cavitated structures in PALLY and PALLY+0.5% DMSO conditions. Scale bar, 50 μm. **c**, Immunofluorescence analysis of the clonal blastoids displays the TE marker (GATA3; cyan), EPI marker (SOX2; yellow), and PE marker (GATA4; magenta). Scale bar, 50 μm. **d**, Graph shows the percentage efficiency of cavitated structures. Data are presented as the mean and standard deviation of three independent experiments. A two-tailed t-test was used, and the P value is as indicated. **e**, Schematic depicting the generation of blastocyst-like structures in 2% DMSO condition. **f**, Representative phase-contrast images showing the cavitated structure in 2% DMSO condition. Scale bar, 50 μm. **g**, Immunofluorescence analysis displays the TE marker (GATA3; cyan), EPI marker (OCT4; yellow), and PE marker (GATA4; magenta). Scale bar, 50 μm. **h**, Graph shows the percentage efficiency of cavitated structures. Data are presented as the mean and standard deviation of three independent experiments. A two-tailed t-test was used, and the P value is as indicated.

We showed that 2% DMSO alone can induce the formation of blastoid (Fig. 4). To investigate whether clonal blastoids could be generated from single nPSCs treated with DMSO alone, we performed our clonal derivation protocol in the presence of 2% DMSO (Fig. 6e). After treatment with 2% DMSO, we successfully obtained cavitated structures from a single nPSC (Fig. 6f). Immunofluorescence analysis revealed the presence of TE- and EPI-like cells (positive for GATA3 and OCT4, respectively), but PE-like cells were absent (Fig. 6g,h), suggesting that future optimizations of the DMSO clonal blastoid protocol (e.g., longer culture or additional factors such as LPA) are required for PE induction.

## DISCUSSION

Understanding the mechanisms of TE development stands at the frontier of research on human blastocysts, implantation, and placental development. Human blastoids based on nPSCs offer a useful model for studying TE specification. Previously, PD^3-9,40^, A83^3-9,40^, and many other small molecules and growth factors have been widely used to drive nPSCs towards the TE fate, a critical step for successful blastoid formation^2,3,8,9^. The use of multiple factors targeting a multitude of gene regulatory and signaling pathways presents significant challenges to understanding of the mechanism of TE differentiation due to pleiotropy and complex crosstalk. Further complicating the matter is the fact that existing blastoid protocols require cell aggregates, which might contain subpopulations that respond to inhibitors and growth factors differently. Here, we show that a single small molecule, DMSO, potently induces TE differentiation from human nPSCs. TE induction by DMSO can be leveraged to enhance blastoid generation and more importantly, enable clonal blastoid generation from single nPSCs without additional pharmacological or genetic perturbations.

DMSO leads to morphological and gene expression changes consistent with those found in TE differentiation induced by PDA83 and TE development in human blastocysts. DMSO also induces a chromatin landscape conducive to TE TF binding. This epigenetic modulation by DMSO provides a deeper understanding of the molecular mechanisms underlying TE differentiation. Blastoids induced by DMSO resemble human blastocysts and the state-of-the-art human blastoid models in morphology, lineage composition, and ability to differentiate into peri-implantation lineages when attached to Matrigel. We show that blastoid formation induced by DMSO requires PKC signaling. DMSO dramatically increases the G1 cell cycle phase of nPSCs by regulating Rb phosphorylation. The G1 phase is important for building up cellular constituents for differentiation, and a longer G1 has been shown to potentiate PSC differentiation^41^, which likely partially explains the differentiation-promoting effect of DMSO. However, the exact mechanism for TE-specific differentiation is still unclear. Future studies may delve deeper into this, potentially uncovering novel insights into human early development.

DMSO, a staple cryoprotectant used in the clinical preservation of human embryos since the 1980s, plays a crucial role in assisted reproductive technologies like in vitro fertilization, where the freezing and storage of embryos are essential for future implantation attempts. Several studies have reported successful births resulting from embryos cryopreserved with DMSO as the cryoprotectant ^42-45^. It is well understood that DMSO aids in cryopreservation by preventing the formation of damaging ice crystals during freezing, thereby ensuring the structural and functional integrity of the embryos ^33^. DMSO is also believed to increase cell membrane permeability by altering membrane dynamics^19^ and facilitate ion transport through transient formation of water pores^46^. The formation of such pores permits the replacement of the intracellular water with cryoprotectants. In cells, membrane effects of DMSO have been proposed to be responsible for its well-known differentiation effect on erythroleukemia cells^21,22^. More interestingly, cell membrane fluctuations^15^ and hydraulic fracturing of cell-cell contacts by fluid buildup^14^ have been shown to regulate the earliest lineage specification events in the embryo. It is tempting to speculate that DMSO may modulate the biophysical properties of nPSC membranes to induce TE differentiation. Future studies using our DMSO model may establish a new paradigm for cell fate specification in totipotent cells.

In cryopreservation protocols, particularly vitrification, higher concentrations of DMSO (∼15% vs. 0.5-2% in this study) are used. Interestingly, several comparative studies showed better survival of human embryos or cells with DMSO-containing cryoprotectants than in DMSO-free ones ^43,47,48^. It is tempting to speculate that remnants of or exposures to DMSO could have contributed to the health of human embryos, similar to what is observed in blastoids.

For the first time, we report a clonal human blastoid model, enabled by the TE-inducing effect of DMSO without additional pharmacological and genetic perturbations, that resembles the human blastocyst. Our future work will focus more on the reproducibility and scalability of clonal blastoids and modeling the human post-implantation process using clonal blastoids. Our clonal blastoids can serve as a valuable in vitro platform for probing fundamental biological inquiries in pre- and post-implantation embryogenesis, modeling conditions associated with early pregnancy, conducting high-throughput drug and genetic screenings, and developing effective contraceptive methods.

## Methods

### Cell culture

The study was reviewed and approved by the KAUST Institutional Biosafety and Bioethics Committee (IBEC). Human PSCs included in this study are chemically reset (cR) cR-SC-9N hiPSCs and cR-H1 hESCs. Primed H1 hPSCs were sourced from the WiCell repository. The RUES2 human ESC line was a generous gift from Dr. Ali H. Brivanlou and was cR in house. These cells underwent epigenetic resetting as per previously established protocol^49^. In summary, primed state PSCs were transitioned into iMEF and treated with PD0325901 (1 μM), LIF (10 ng/mL), and VPA (1 mM) for 3 days. Subsequently, cells were transferred to a medium containing 1 μM PD0325901, 2 μM XAV-939, 2 μM Gö6983, and 10 ng/mL LIF (PXGL). By day 10, naïve dome-shaped colonies emerged and were purified through FACS sorting for SUSD2 or via gelatinization for several passages. Cell culture was maintained at 37°C under 5% CO2 and 5% O2.

### Trophectoderm and blastoid formations

To prepare PXGL nPSCs for trophectoderm and blastoid differentiation, nPSCs are grown on iMEF in PXGL medium for 3-4 days. Next, cells are dissociated using TrypLE for 5 min at 37°C with gentle pipetting until a single-cell suspension is achieved. TrypLE is then inactivated using 0.2% BSA N2B27 medium, and cells are pelleted by centrifugation. To remove any remaining iMEF, cells are seeded on a 0.1% gelatin-coated surface for 60 min. Following this, cells are passed through a 40 µm strainer to eliminate clumps, centrifuged once more, and resuspended in an appropriate medium. For trophectoderm differentiation, a previously established protocol was followed^1^. Briefly, cells are resuspended in PXGL medium containing ROCK inhibitor and plated onto laminin-coated dishes. After 24 h, the medium was replaced with N2B27 supplemented with PD03 (1 μM) and A83 (1 μM) or Retinoic acid (varying concentrations) for 96 h, after which the cells were harvested for downstream analysis. Cells remain in this medium for 3 – 4 days with daily medium exchanges. DMSO was used as a vehicle for PD03 and A83 inhibitors, and RA. The effective DMSO concentration in the TE experiments was 0.004%, 0.004%, and 0.01% for PD03, A83 and RA, respectively. For blastoid formation, we have followed an established protocol^6^. Cells are resuspended in N2B27 medium supplemented with ROCK inhibitor and seeded at a density of approximately 75 cells per microwell in 400 µm AggreWell plates (Stem Cell Technology, cat# 34425). Once cell structures appear solid, the medium is changed to N2B27 supplemented with PALLY (1 μM PD03, 1 μM A83, 500 nM LPA, 10 ng/mL LIF, and 10 μM ROCK inhibitor) and maintained for two days or until cavitation is observed. Lastly, the medium is changed to N2B27 supplemented with LPA (500 nM) and ROCK inhibitor (10 µM) (LY) for an additional two days to allow the structures to mature. DMSO was used as a vehicle for PD03 and A83 inhibitors. For the blastoids grown in PALLY, the overall effective DMSO concentration during the PALLY phase was 0.008%, which is much lower than the concentrations of DMSO that have effects on blastoid formation. The following morphometric criteria were followed to describe the blastoids. Blastoids are defined as a single-layer cavitated structure with a diameter between 150-250 µm and the presence of a singular inner cell mass. Non-cavitated or multi-cavitated structures were excluded from downstream analyses.

### Clonal blastoid derivation from a single nPSC

To generate single-cell clonal blastoids from SC-9N nPSCs, we followed a standard protocol using the Aggrewell 400 plate (24-well format). The nPSCs were cultured in the PXGL medium, with daily media replacement and passage every three days to maintain their healthy dome-shaped morphology. Prior to seeding, we added the ROCK inhibitor to the PXGL medium and introduced it to the cells one day in advance. On the day of seeding (referred to as day 0), we harvested nPSCs using TrypLE and DNase I. We confirmed the successful digestion of the cells into single cells through microscopic examination. It is worth noting that DNase I concentration and incubation time for digestion can be adjusted as needed to achieve single-cell status. Subsequently, we seeded the nPSCs at a density ranging from 700 to 900 cells per well in N2B27 medium containing the ROCK inhibitor. The exact seeding density may vary based on cell type, but we visually confirmed the single-cell seeding under the microscope. To enrich for single cells, we can also perform FACS sorting on the pooled population. Next, we treated the nPSCs with N2B27 medium supplemented with the ROCK inhibitor alternate days until compacted aggregate formation was observed, typically within 6-8 days. Once the compacted aggregate was formed, we exposed the structures to the following treatment conditions: PALLY alone (as shown in Fig. 3a), PALLY supplemented with 0.5% DMSO (as shown in Fig. 3a), or 2% DMSO (as shown in Fig. 4a), for a duration of 48 h. Subsequently, for the former two conditions, the structures received treatment with LPA and the ROCK inhibitor, while the latter condition was treated with the ROCK inhibitor alone for an additional 48 h. After completing the final treatment phase, we imaged the structures and harvested them for downstream analyses.

### Immunofluorescence

All samples, except for blastoids, were grown on ibidi 8-well chambers (cat# 80826). The samples were washed with PBS and fixed with 4% PFA for 15 min. Afterward, the samples were washed three times with PBS and permeabilized and blocked using a solution containing 0.2% Triton-X-100 and 6% normal donkey serum for 1 h. The same solution was used to dilute the primary antibody according to the manufacturer’s recommendations and incubate the samples overnight. Following incubation, the samples were washed three times with a solution containing 0.2% Triton-X-100 and 6% normal donkey serum. The secondary antibodies were diluted at a 1:500 ratio and used to incubate the samples for 1 h at 37°C. The samples were washed three more times with PBS. Samples other than blastoids were mounted using Prolong Antifade solution with DAPI, while blastoids were kept in PBS and DAPI. Finally, images were acquired using a Leica SP8 or Stellaris 8 confocal microscope and processed using the Fiji software.

### Flow cytometry

Samples were dissociated by incubating in TrypLE for 5-15 min with agitations and were run through a 40 μm strainer. Then, samples were stained with FACS-grade antibodies in a FACS buffer solution for 30 min. For cell cycle analysis, samples were fixed in pre-cold 80% ethanol at ™20°C overnight. Samples were then washed with PBS and incubated in 200 μg/mL RNase A at 37°C for 30 min. Each sample was stained with 10 μL propidium iodide (PI) staining solution (Tonbo Biosciences, cat# 13-6990-T200) for 20 min. All samples were analyzed and/or with BDFACSAria Fusion. Data was processed in the BD FlowJo software.

### Bulk ATACseq and RNAseq

RNA was extracted using QAIGEN RNAeasy kit (cat#74104) and quality-checked with qubit and bioanalyzer 2100. RNA was outsourced to Novogene for further QC, stranded library construction, and sequencing with an output of 9 Gb raw data. Moreover, ATACseq library construction was performed using a commercial kit, Diagenode (cat#C01080002) and in accordance with the manufacturer’s recommendations. Ready to sequence libraries were sequenced by Novogene Inc.

### Bulk RNAseq data analysis

Quality control, alignment, gene annotation and read count quantification were performed by Sequentia Inc. For public RNA-seq datasets, we downloaded the raw data and performed de novo analysis to obtain the raw counts of the samples using the same pipeline by Sequantia. The batch effects of the samples were removed using ComBat. For global analyses, we considered only genes with a count of 15 or higher in at least two samples. Principal component, differential expression and cluster analyses were performed based on log2 expression values computed with custom scripts, in addition to the Bioconductor packages DEseq2. For PCA of the RNA-seq data, plotPCA() in the DEseq2 package was used to normalize the counts and compute the PCA data. PCA data was plotted with the ggplot2 package in R (http://ggplot2.org). Gene density that contributed to PCA plots were calculated using kernel density estimation. Heatmaps were plotted by the pheatmap package in R using Z-scores. Log2 expression values were used to plot dotplots of marker expression.

### Bulk ATAC-seq data analysis

The ATAC-seq sequencing data were preprocessed with the default parameters by NGmerge to remove adapters and low-quality reads. The cleaned reads were aligned to the human genome assembly (hg38) using bowtie2 (v.2.3.4.1) with the default parameters except for the following options: “-X 2000 --no-unal --very-sensitive”. Reads mapping to mitochondrial DNA were discarded using the “grep –v chrM” command. For downstream analysis, PCA duplicates were removed using the sambamba (v. 0.6.7-pre1) markdup function with “-r” parameters. Alignment BAM files were transformed into read coverage files (bigWig format) using deepTools. The hg38 blacklist regions were removed using “--blackListFileName” parameters. Peaks were called using MACS2 (v.2.1.1.20160309) with default options except for the following options: “--nomode-f BAMPE --keep-dup all”. ATAC-seq peaks across all samples were merged into one union ATAC-seq peak set using the BEDTools (v.2.26.0) merge function, and ATAC-seq reads in each sample were calculated over the union ATAC-seq peak set using the deepTools (v. 3.5.1) multiBigwigSummary function with bigWig files. Peak annotation of the union ATAC-seq peak set was performed by chIPseeker. The output matrix was then log2 transformed and used as input for the PCA. The PCA plot and heatmap were then plotted with the ggplot2 package in R using the normalized ATAC-seq reads over each peak of the growth and differentiation marker genes. The prediction of transcription factor binding sites with footprints and motif enrichment were performed by HINT-ATAC.

### Single-cell RNA-seq library construction and sequencing

We collected pre-implantation blastoid structures and pooled them into a 10 cm dish containing N2B27 media with ROCK inhibitor. We selected 150-250 μm-sized blastoids under a dissection microscope, washed them with PBS, and transferred them to a 1.5 mL Eppendorf tube containing TrypLE and DNase I. These structures were then placed in a 37°C mixer and agitated at 800 RPM for 15 min. Following this, the dissociated blastoids were passed through a 40 µm strainer and stained with a TROP2 trophectoderm marker. To capture a sufficient number of primitive endoderm cells, we employed FACS sorting to dilute TROP2-positive cells. We then utilized the 10X Chromium Single Cell 3’ Solution V2 for library preparation, targeting 5000 cells per reaction and adhering to the manufacturer’s protocol. Similarly, we digested post-implantation blastoids with TrypLE, counted the cells, and proceeded with single-cell library preparation. Ready-to-sequence libraries were sent to Novogene for sequencing.

### Single cell RNA-seq data analysis

Raw FASTQ files underwent standard quality control (QC) procedures with FastQC v0.11.9 and BBDuk v35.85, trimming low-quality bases, removing adapter sequences, and filtering low-quality reads with set parameters of a minimum 35 bp read length and a quality score of 25.

Using STARsolo, the high-quality reads were mapped against the reference human genome (GRCh38/Ensembl release 104), resulting in matrices of raw counts. An additional QC in R filtered out cells with minimal counts across genes and those with high mitochondrial gene expression percentages. The matrices were normalized using the scaling normalization technique in the Batchelor R package. The Scran R package facilitated dimensionality reduction, graph-based cell clustering, cluster visualization, and cluster marker gene identification. Clusters were annotated based on recognized publications and curated lists of cell type marker genes.

Doublet detection incorporated two strategies from the scDblFinder R package (https://bioconductor.org/packages/release/bioc/html/scDblFinder.html). The initial method identified doublets as clusters situated between two distinct clusters in terms of expression profiles. The secondary approach involved simulating doublets from expression data and employing a classifier to pinpoint doublet cells.

Following previously reported guidelines^6^, dataset integration was achieved. Published datasets^6,27-29^ were downloaded, processed, and transformed into count expression matrices. These matrices underwent filtration steps, considering parameters like mitochondrial counts and gene counts. Subsequent processes, such as data normalization and integration, combined functions and tools from the Scran, Batchelor, and SeuratWrappers R packages. The results included graph-based clustering, formed on the MNN low-dimensional coordinates, and a UMAP visualization designed from the leading 20 principal components. The identification of cluster markers and annotations was consistent with previously mentioned techniques.

Enrichment analyses leveraged the Kyoto Encyclopedia of Genes and Genomes (KEGG) and GO databases, examining the marker gene lists for each cluster. Pathways and terms were evaluated through hypergeometric tests, subjected to FDR corrections, and significant findings were highlighted. Lastly, the pathview package v1.30.1 visually represented gene expression within specific functional pathways.

### Real-time quantitative PCR

Total RNA was extracted using a Qiagen kit (cat# 74104) following the manufacturer’s protocol and quality checked with Nanodrop. Using 1 μg of this RNA, reverse transcription to cDNA was carried out with the Bio-Rad iScript cDNA synthesis kit (cat#1708890) as per manufacturer recommendations. Subsequent qPCR was performed on the Bio-Rad CFX384 detection system using Bio-Rad Ssoadvanced SYBR polymerase (cat#1725270). Data analysis is performed in Excel using ΔΔCt method relative to GAPDH. All Oligos sequences used in this study are listed in Supplementary table 3.

### Western blot

Cells were lysed using RIPA Lysis Buffer (150 mM NaCl, 1.0% Triton X-100, 0.5% Na-deoxycholate, and 50 mM Tris pH 8.0) with protease and phosphatase inhibitor cocktails (Sigma, cat#11836170001, and Life Technologies, cat# 78440, respectively). The samples were prepared using 2x Laemmli buffer (Bio-rad, cat#1610747) according to the manufacturer’s instructions, and 10 µg/sample was separated on 4-12% Bis-Tris Plus Gels (ThermoFisher, cat#NW04125BOX) and transferred to nitrocellulose membranes. The membranes were blocked in 5% skim milk or BSA in 0.1% TBS-Tween® 20 following antibody instructions for 1 h at room temperature and then incubated overnight at 4°C with the primary antibodies. After washing, the membranes were incubated with secondary antibodies for 1 h at room temperature, then incubated in Chemiluminescent HRP substrate (ThermoFisher, cat# 34076) for signal detection. Antibodies details are described in the Supplementary table 1. Protein expression analysis was conducted with ImageJ software.

### Blastoids attachment assay

Blastoid in vitro implantation was executed based on prior studies^2,50,51^. By day 5, high quality blastoids were manually selected under a dissection microscope. These were then settled in a Matrigel-pre-coated well for 1 h, immersed in IVC1 media. This medium comprised Advanced DMEM/F12 (Thermo Fisher Scientific, cat#12634-010), 20% Heat-inactivated FBS (Thermo Fisher Scientific, cat#30044333), 2 mM L-GlutaMAX (cat#35050-061), 0.5% penicillin-streptomycin mix (cat#15140-122), 1% ITS-X (cat#51500-056), and 1% sodium pyruvate. It also included 8 nM β-estradiol (Sigma-Aldrich, cat#E8875), 200 ng/mL progesterone (cat#P0130), 25 μM N-acetyl-L-cysteine (cat#A7250), and Y27632 (Selleckchem, cat#S1049). After two days in IVC1, the medium was swapped to IVC2, mirroring IVC1’s composition but with a 20% FBS to 30% knockout replacement serum shift. Following four days, the medium was retained for hCGB assay, and implanted blastoids were dissociated into individual cells using TrypLE for subsequent experiments.

### PKC inhibitor assay

For the Pan PKC inhibitor assay, 1 μM of Gö6983 (Selleckchem) was added during the compaction stage under the mentioned conditions. The structures were collected on day 6 and subjected to immunofluorescence analysis to check the expression of GATA3, YAP1, and OCT4. For isozyme-specific peptide inhibitors against PKCα/ý, PKCý, and PKCη isoforms, 100 nM of Gö6976 (Cell Signaling), 10 μM of PKCý pseudosubstrate inhibitor (Life Technologies), and 10 μM of PKCη pseudosubstrate inhibitor (Millipore) were treated during the compaction stage and cavity formation was examined under the microscope.

### Statistical analysis and reproducibility

No statistical methods were employed to pre-determine the sample size. The experiments were conducted without randomization. Furthermore, the investigators were not blinded to the allocation during both the experiments and the outcome assessment. Results were documented as means and standard deviations from at least three independent experiments (unless otherwise specified in the respective figure legends or within the figures). Comparisons between the two sets were analyzed using the Student’s t-test. Experiments involving three or more groups were examined by one-way or two-way analyses of variance followed by the mentioned post hoc test. A P-value <0.05 was considered statistically significant. All of the statistical analyses were performed using GraphPad Prism 10 software.

### Data availability

scRNA-seq, bulk RNA-seq, and bulk ATAC-seq were deposited at the Gene Expression Omnibus (GEO) repository. The accession numbers are as follows; scRNAseq datasets (GSE237669), bulk RNA-seq (GSE236961), and bulk ATAC-seq (GSE239744).

## Acknowledgments

This work was supported by KAUST Office of Sponsored Research (OSR), under award number BAS/1/1080-01. We thank the members of the Li lab, Jinna Xu, Chongwei Bi, Mengge Wang, Xuan Zhou, Yeteng Tian, and KAUST Imaging and Characterization Core Lab and Bioscience Core Lab for their help in this research.

## Author contributions

S.A. performed the experiments related to the DMSO triggering trophectoderm differentiation and blastoid formation; A.P.C. performed experiments related to DMSO-only blastoid formation, mechanism-related studies, and clonal derivation of blastoids; Y.J. performed experiments related to 2D trophectoderm differentiation and cell cycle analysis. Y.Z. and A.V. performed the bioinformatic analysis. Y.J., I.S., and G.R.M. assisted S.A. and A.P.C in performing experiments. B.Y. assisted S.A. in scRNAseq analysis; A.P.C., S.A., A.V., and M.L. wrote the manuscript. J.C.I. and M.L. conceived and supervised the study.

## Supplementary movie legend

**Supplementary movies 1 and 2**: Live cell image analysis shows the formation of blastoids from nPSCs under PALLY (supplementary movie 1) and PALLY+0.5% DMSO (supplementary movie 2) conditions.

## Supplementary Figure legends

**Supplementary Fig. S1.**
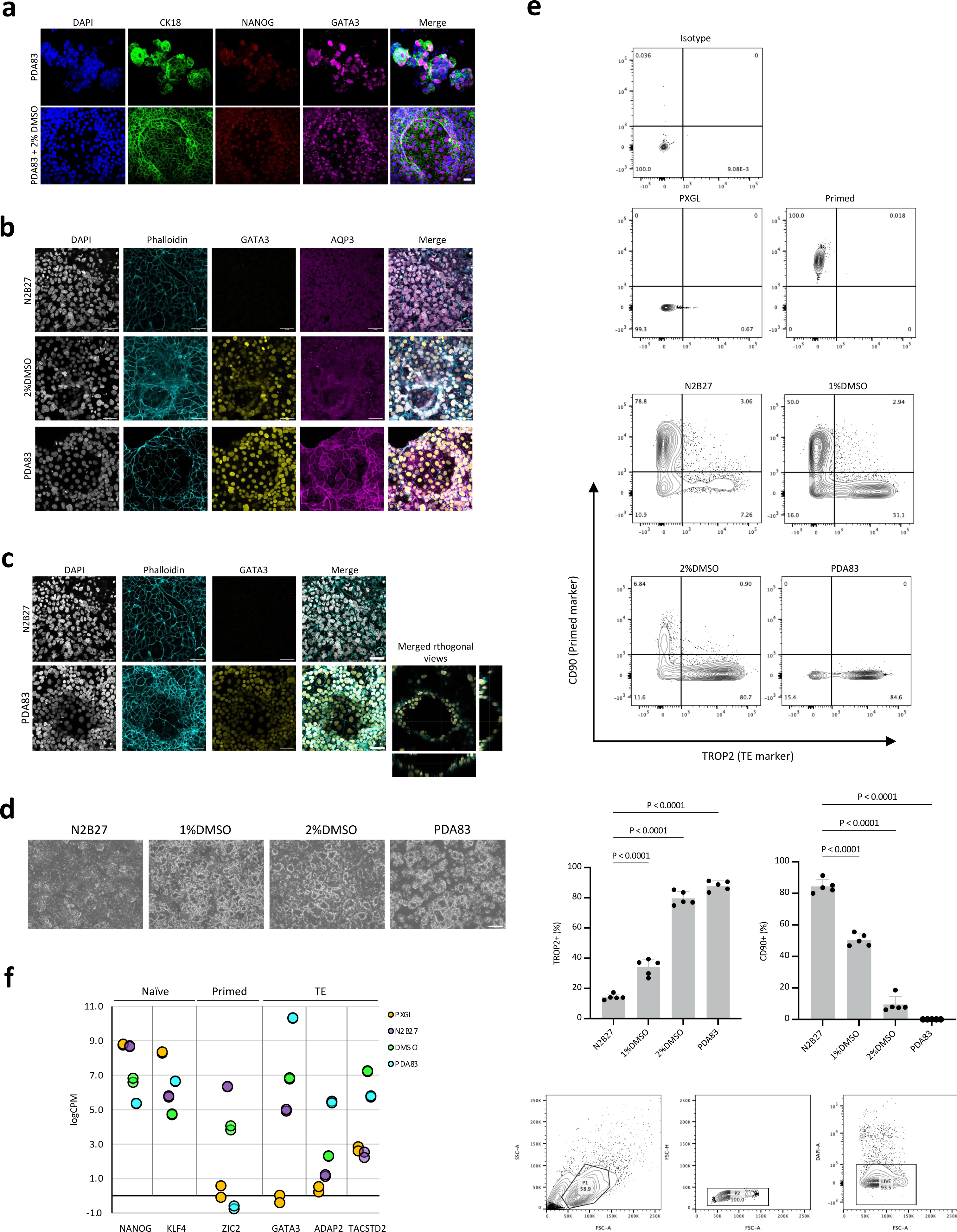
DMSO promotes exit of naïve state and facilitates TE differentiation. **a**, Immunofluorescence analysis shows the expression of CK18 (green), NANOG (red), and GATA3 (magenta) in PDA83 and PDA83+2% DMSO conditions. Scale bar, 40 μm. **b**, Immunofluorescence analysis shows the expression of Phalloidin (cyan), GATA3 (yellow), and AQP3 (magenta) in N2B27, 2% DMSO, and PDA83 conditions (n = 3). Scale bar, 50 μm. **c**, Immunofluorescence analysis shows the expression of Phalloidin (cyan) and GATA3 (yellow) in N2B27 and PDA83 conditions. Scale bar, 50 μm. **d**, Representative brightfield images display the cyst formation under different conditions. Scale bar, 400 μm. **e**, FACS data displays the population of cells in TROP2 (a marker for TE) and CD90 (a marker for primed state) in different conditions (n = 3). **e**, qRT-PCR data shows the expression pattern of various genes in naïve, primed, and TE states under different culture conditions (n = 2) (**top**), graphs show the quantification of TROP2 and CD90 populations (**middle**), and its gating strategy (**bottom**).

**Supplementary Fig. S2.**
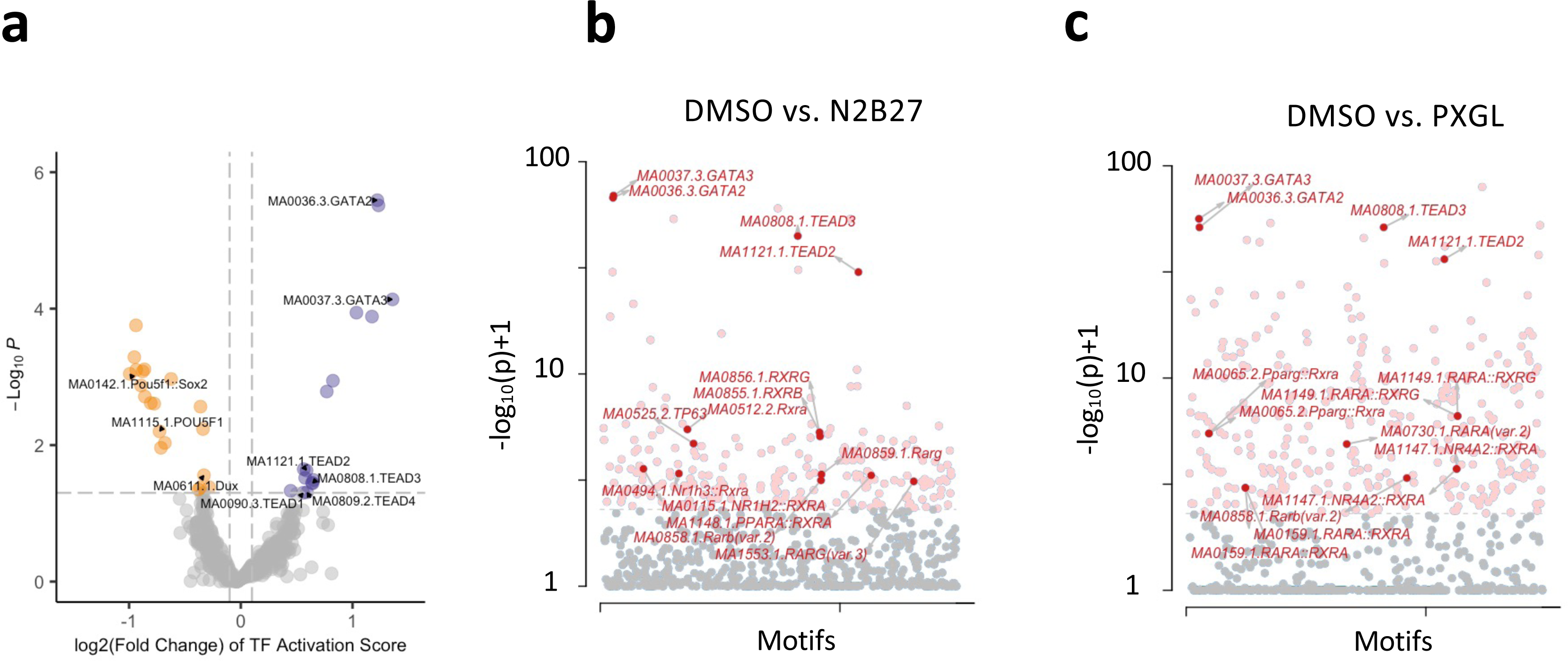
Motif enrichment analysis. **a-c.** Motif enrichment analysis shows the effect of DMSO on TE transcriptional factors (**a**), DMSO vs. N2B27 (**b**) and DMSO vs. PXGL (**c**).

**Supplementary Fig. S3.**
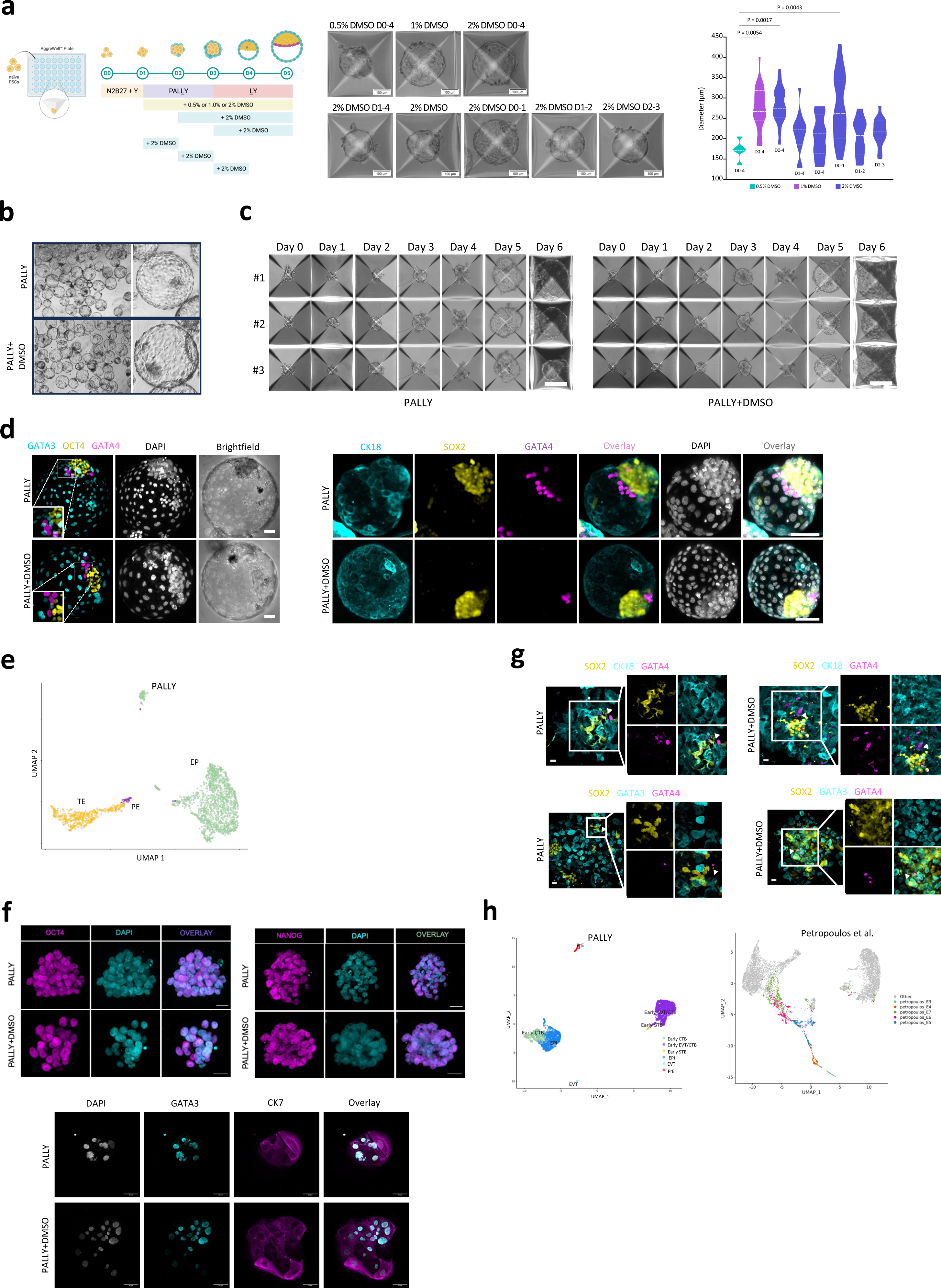
DMSO improves the quality of the PALLY-derived blastoids. **a**, Scheme explains the design of the dose and time window of DMSO exposure in PALLY protocol (**left**). Brightfield images show the formation of blastocyst-like structures under different conditions. Scale bar, 100 μm (**middle**). The violin plot shows the diameter (**right**). **b**, Brightfield images showing the blastocyst-like structures formed in PALLY and PALLY+0.5% DMSO conditions. Scale bar, 200 μm. **c**, Brightfield images showing the blastocyst-like structures in PALLY and PALLY+0.5% DMSO conditions, consistently. Scale bar, 200 μm. **d**, Immunofluorescence analysis shows the expression of GATA3 (cyan), OCT4 (yellow), and GATA4 (magenta) (left panels) and the expression of CK18 (cyan), SOX2 (yellow), and GATA4 (magenta) (right panels). Scale bar, 60 μm. **e**, UMAP of the transcriptome of 4687 single cells of the pre-implantation PALLY blastoids with cell type annotation. **f**, Immunofluorescence analysis shows the rederivation of stem cells (**top left**, OCT4; **top right**, NANOG; and **bottom**, GATA3/CK7) in PALLY and PALLY+0.5% DMSO conditions. Scale bar, 20 μm. **g**, Immunofluorescence analysis of post-implantation structures shows the expression of SOX2 (yellow), CK18/GATA3 (cyan), and GATA4 (magenta) in PALLY+0.5% DMSO condition. Scale bar, 20 μm. **h**, UMAP of the transcriptome of 3875 single cells of the attached PALLY blastoids with cell type annotation (left). UMAP projects of integrated datasets showing cells from published Petropoulos et al. (right).

**Supplementary Fig. S4.**
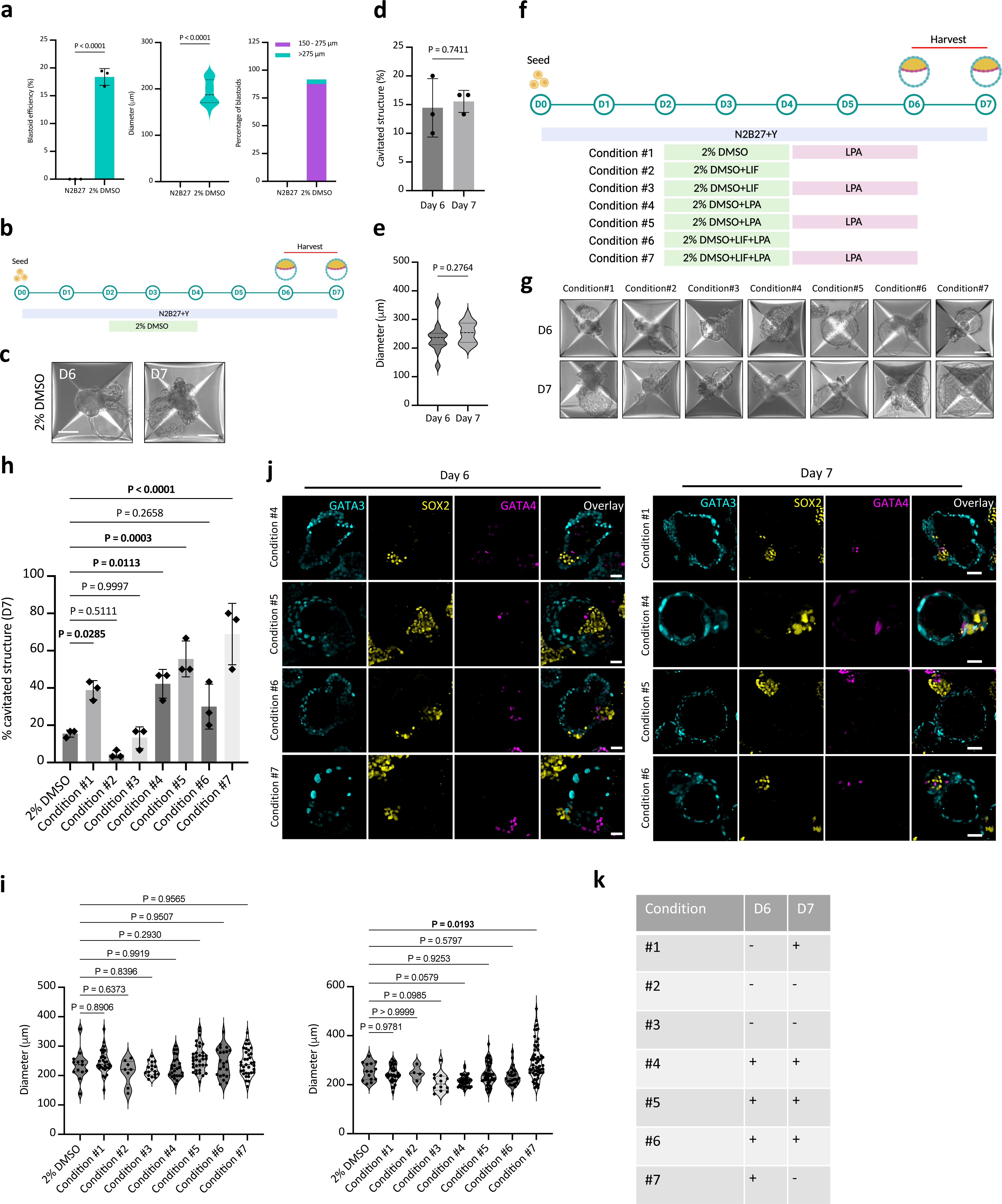
LPA is indispensable for the DMSO-derived blastoid generation. **a**, Percentage of cavitated structures (**left**), size (**middle**), and percentage size distribution (**right**) of the structures in 2% DMSO condition. Data are presented as the mean and standard deviation of three independent experiments. A two-tailed t-test was used, and the P values are as indicated. The morphometric definition of human blastocyst-like structures is defined in *Methods*. **b**, Schematic representation of the extended culture of human blastocyst-like structures in DMSO condition. **c**, Representative phase-contrast images of DMSO-derived cavitated structures harvested from the microwells on D6 and D7. Scale bar, 100 μm. **d**,**e**, Comparison of percentage cavitation (**d**) and size (**e**) of the D6 and D7 structures. Data are presented as the mean and standard deviation of three independent experiments. A two-tailed t-test was used, and the P values are as indicated. **f**, Schematic representation of the generation of human blastocyst-like structures using various conditions. **g**, Representative phase-contrast images of cavitated structures harvested from the microwells using various treatment conditions and time points. Scale bar, 100 μm. **h**, Graph shows the percentage cavitation in different treatment conditions in D7 structures. Data are presented as the mean and standard deviation of three independent experiments. One-way ANOVA followed by the Dunnett post hoc test was used, and the P values are as indicated. **i**, Graphs show the diameter of the cavitated structures in different treatment conditions in D6 (**left**) and D7 (**right**) structures. Data are presented as the mean and standard deviation of three independent experiments. One-way ANOVA followed by the Dunnett post hoc test was used, and the P values are as indicated. **j**, Immunofluorescence analysis of the TE marker (GATA3; cyan), EPI marker (SOX2; yellow), and PE marker (GATA4; magenta) in various treatment conditions (D7). Scale bar, 50 μm. **k**, Summary table shows the effect of various treatment conditions and time window on trilineage development.

**Supplementary Fig. S5.**
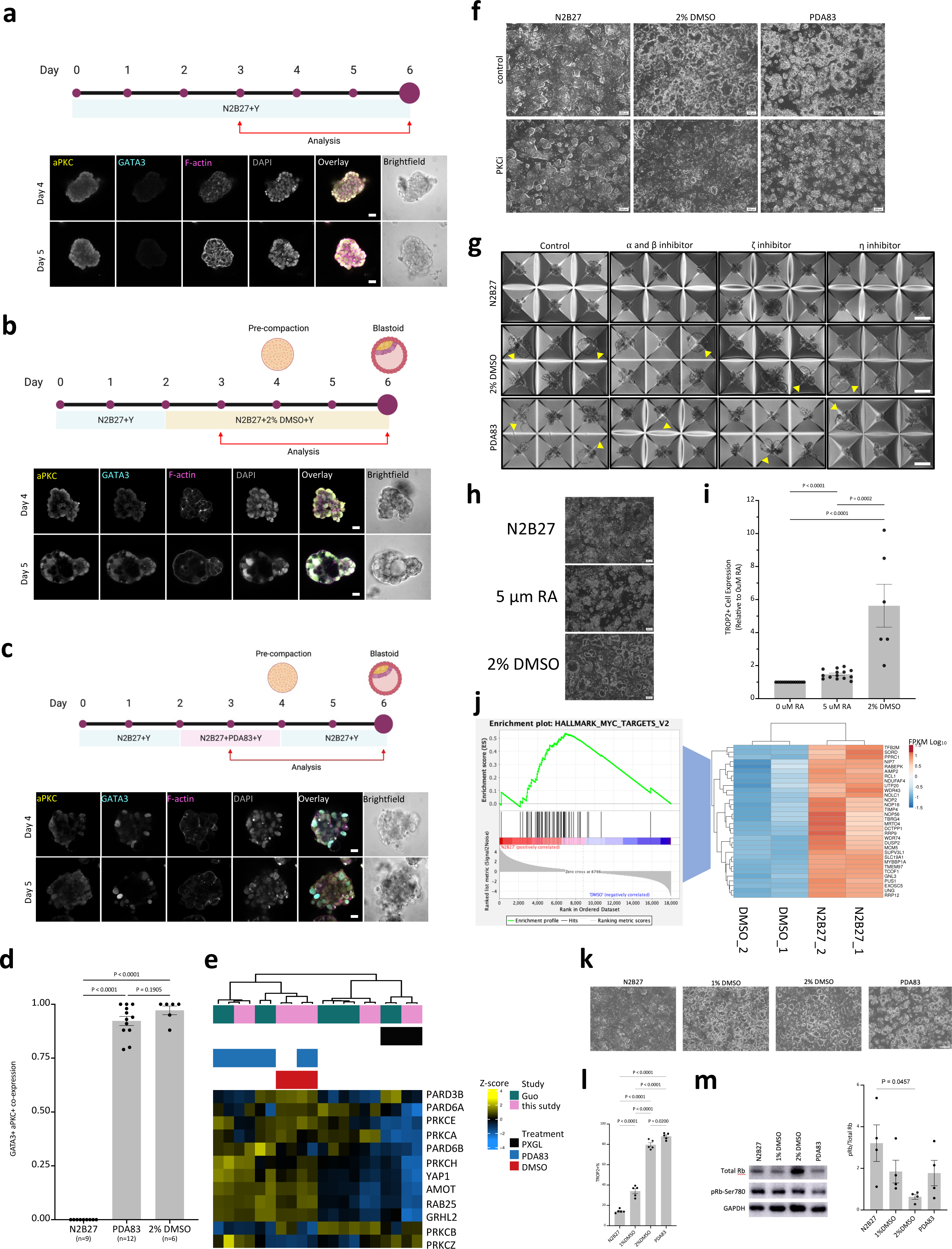
PKC mediates the cavity formation in DMSO-derived blastoids. **a-c**, Immunofluorescence analysis shows the colocalization pattern of aPKC (yellow), GATA3 (cyan), and F-actin (magenta) in N2B27 (**a**), 2% DMSO (**b**), and PDA83 (**c**) conditions. Scale bar, 25 μm. Schematic representation of the treatment process is depicted in respective conditions. **d**, Quantitative analysis of day 6 structures from (Fig. 5a). Data are presented as the mean and standard deviation. The number of structures analyzed is as indicated. One-way ANOVA followed by Tukey’s post hoc test was used, and the P values are as indicated. **e**, Heatmap shows the enrichment of aPKC cell polarity genes. **f**, Representative brightfield images of day 4 samples in N2B27, 2% DMSO, and PDA83 conditions with and without treatment with PKC inhibitor. **g**, Representative brightfield images of day 6 structures in N2B27, 2% DMSO, and PDA83 conditions (n = 3). The yellow arrow indicates proper cavitated structures. Scale bar, 200 μm. **h**, Representative brightfield images of day 4 samples in N2B27, 2% DMSO, and RA conditions (n = 3). Scale bar, 200 μm. **i**, FACS analysis of samples from (**h**). Data are presented as the mean and standard deviation (control; n = 14, 5 μm RA; n = 14, and 2% DMSO; n = 6). One-way ANOVA followed by Tukey’s post hoc test was used, and the P values are as indicated. **j**, Heatmap shows the gene expression in DMSO and N2B27 conditions. **k**, Representative brightfield images of day 4 samples in N2B27, 1%/2% DMSO, and RA conditions (n = 3). Scale bar, 200 μm. **l**, FACS analysis of samples from (**k**). Data are presented as the mean and standard deviation of three independent experiments. One-way ANOVA followed by Tukey’s post hoc test was used, and the P values are as indicated. **m**, Western blot analysis shows the expression of the mentioned proteins (**left**), and quantification shows the normalized pRb-Ser780 expression (**right**). GAPDH was used as a loading control. Data are presented as the mean and standard deviation (n = 4). One-way ANOVA followed by Tukey’s post hoc test was used, and the P values are as indicated.

**Supplementary Fig. S6.**
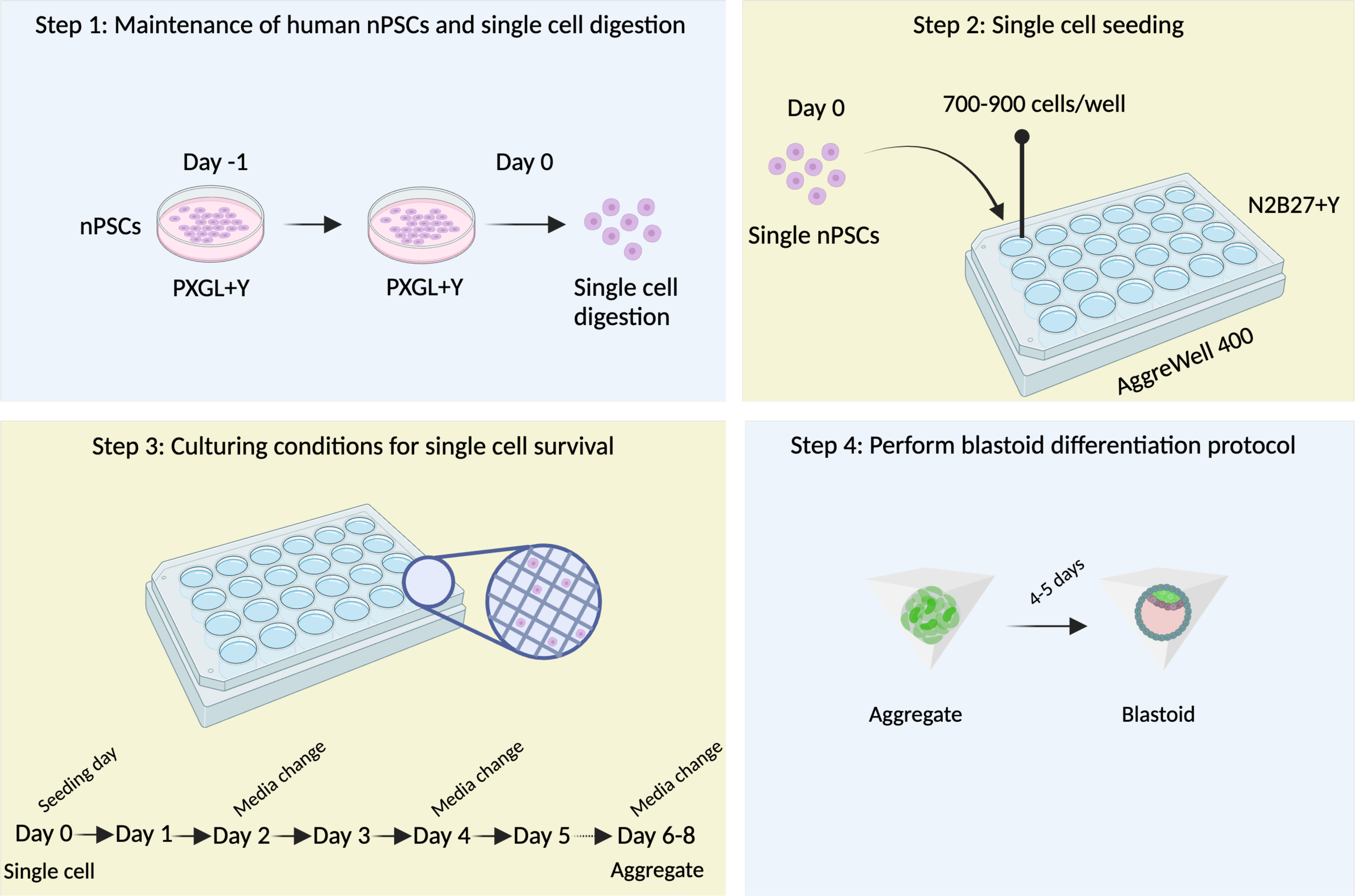
Schematic representation shows the derivation of clonal blastoids from a single nPSC.

## Supplementary tables

**Supplementary table 1:** Antibody list used for immunofluorescence and western blot.

| Antibody | Vendor | Catalog number | Dilution |
| --- | --- | --- | --- |
| CK7 | Abcam | Ab181598 | 1:100 |
| CK18 | Santa Cruz | sc-31700 | 1:50 |
| OCT3/4 | Santa Cruz | Sc-9081 | 1:50 |
| NANOG | Abcam | Ab77095 | 1:100 |
| YAP1 | Abcam | Ab205270 | 1:200 |
| GATA4 | Cell signaling | MAB36966S | 1:400 |
| PODXL | Cell signaling | MAB55601 | 1:200 |
| SOX2 | Cell signaling | MAB4900S | 1:200 |
| HLA-G | Cell signaling | MAB79769 | 1:100 |
| Syncytin1 | Bioss | BS-2962R | 1:100 |
| AQP3 | Invitrogen | PA5-77840 | 1:200 |
| GATA3 | R&D Systems | MAB6300 | 1:200 |
| GATA3 | Cell signaling | MAB5852S | 1:200 |
| PKCzeta | Abcam | Ab59364 | 1:200 |
| RB(4H1) | Cell signaling | MAB14731S | 1:1500 |
| Phospho-RB | Cell signaling | MAB3590 | 1:2500 |

**Supplementary table 2:**
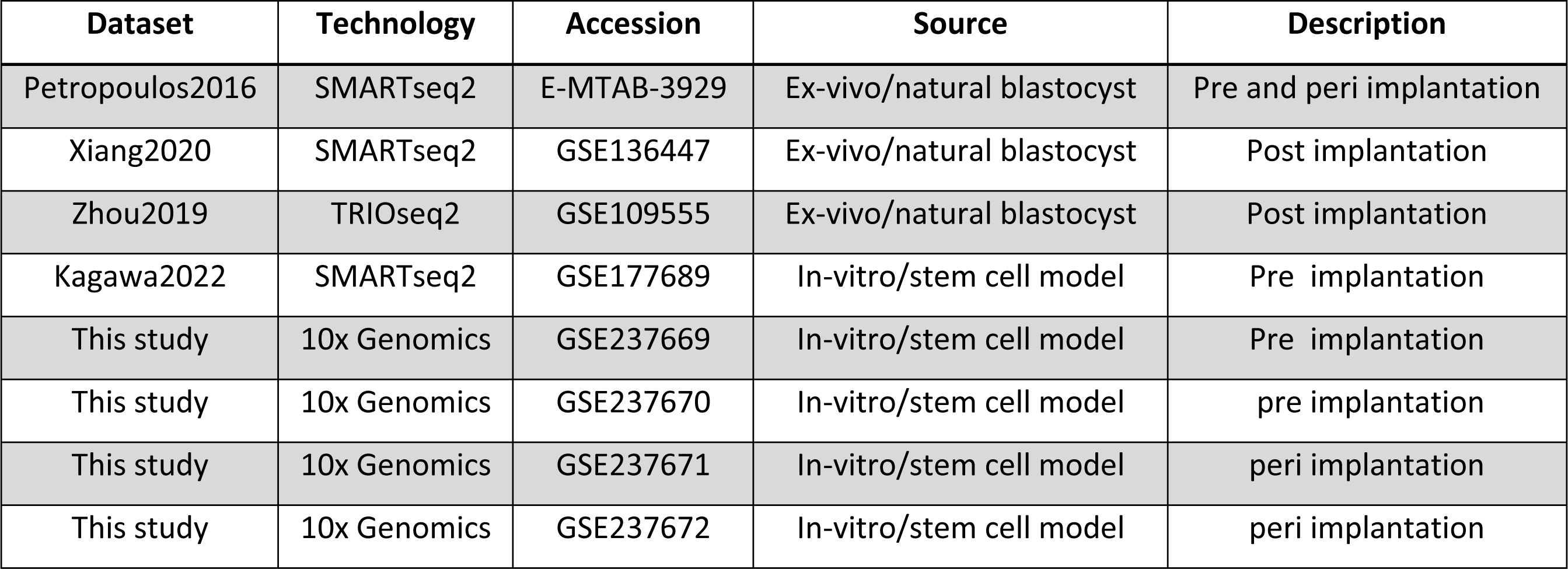
scRNAseq datasets descriptions used and produced in this study.

| Dataset | Technology | Accession | Source | Description |
| --- | --- | --- | --- | --- |
| Petropoulos2016 | SMARTseq2 | E-MTAB-3929 | Ex-vivo/natural blastocyst | Pre and peri implantation |
| Xiang2020 | SMARTseq2 | GSE136447 | Ex-vivo/natural blastocyst | Post implantation |
| Zhou2019 | TRIOseq2 | GSE109555 | Ex-vivo/natural blastocyst | Post implantation |
| Kagawa2022 | SMARTseq2 | GSE177689 | In-vitro/stem cell model | Pre implantation |
| This study | 10x Genomics | GSE237669 | In-vitro/stem cell model | Pre implantation |
| This study | 10x Genomics | GSE237670 | In-vitro/stem cell model | pre implantation |
| This study | 10x Genomics | GSE237671 | In-vitro/stem cell model | peri implantation |
| This study | 10x Genomics | GSE237672 | In-vitro/stem cell model | peri implantation |

**Supplementary table 3:**
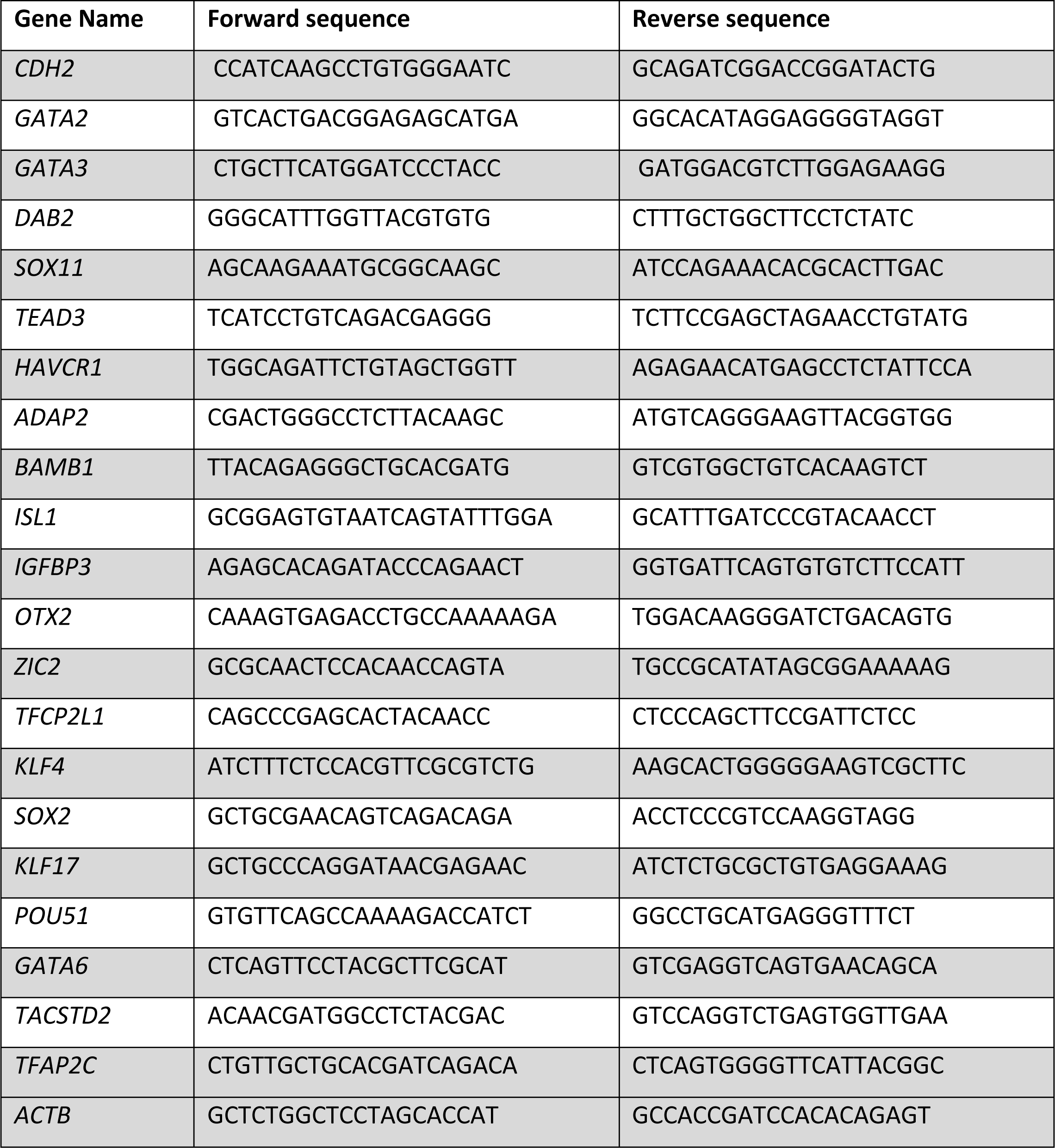
Oligos list used in this study for real-time quantitative PCR.

